# B cell responses specific for polyomavirus-derived oncoprotein are predictive of Merkel cell carcinoma tumor control

**DOI:** 10.1101/2025.07.22.666002

**Authors:** Haroldo J. Rodriguez, Allison J. Remington, Matthew D. Gray, Rian Alam, Macy W. Gilmour, Carina Morningstar, Gabriel F. Alencar, Thomas Pulliam, Erin McClure, Neha Singh, Francesca Urselli, Katrina Poljakov, Kimberly Smythe, Rima Kulikauskas, Kristin L. Robinson, Ata S. Moshiri, Cecilia C.S. Yeung, Minggang Lin, Kristen Shimp, Allison Schwartz, Anne M. Macy, Marti R. Tooley, Melissa L. Baker, Joseph J. Carter, Kayla Hopwo, Naina Singhi, Jakob Bakhtiari, Mikel Ruterbusch, Carolyn Shasha, Maria Iuliano, Logan J. Mullen, Blair L. DeBuysscher, Joshua R. Veatch, David M. Koelle, Denise A. Galloway, Paul Nghiem, Justin J. Taylor

## Abstract

Merkel cell carcinomas typically arise from clonal integration of the Merkel cell polyomavirus. Immunogenic viral oncoproteins then lead to tumorigenesis. Oncoprotein-specific T cells are essential for anti-MCC immunity, but it is unclear whether B cells promote tumor control. Here, we analyzed the frequency and phenotype of viral oncoprotein-specific and total B cells in 47 blood samples and 19 unmatched tumors from MCC patients— of which 8 out 19 progressed. The phenotype of blood B cells did not correlate with MCC patient outcomes. In contrast, all 11 patients with robust oncoprotein-specific antibody-secreting and/or germinal center B cells in tumors experienced long-term MCC control. *In vitro*, B cells engineered to be specific for viral oncoproteins increased the sensitivity of oncoprotein-specific CD4+ T cells by over 50-fold. Together, our findings suggest that cancer-specific B cells promote anti-tumor immunity via increased T cell responses and that cancer-specific B cell augmentation could be therapeutically relevant.

**Statement of Significance:** The link between cancer-specific B cells in anti-tumor immunity and clinical outcomes remains poorly defined. Here, we show that tumor-associated B cells specific for a viral oncoprotein expressed in MCC patient tumors predict disease control with remarkable accuracy, establishing their potential as active participants in tumor immunity.

## Introduction

Merkel cell carcinoma (MCC) is a rare aggressive skin cancer with a mortality rate of ∼30%^1–4^. MCC is usually driven by integration of Merkel cell polyomavirus (MCPyV) T-antigen (T-Ag) DNA into host chromosomes. This leads to constitutive expression of the viral small and truncated large T-Ag oncogenic proteins responsible for tumorigenesis^5–7^. T-Ag oncoproteins are immunogenic and targeted by T cells^8–11^ and B cells^12,13^. While studies have established the importance of MCPyV T-Ag-specific CD8^+^ T cells in MCC tumor control, the role that antibodies and B cells play in anti-tumor immunity is unknown.

Most patients with T-Ag-driven MCC have T-Ag-specific antibodies in blood that are largely undetectable in individuals without MCC^12,13^. The level of T-Ag-specific antibodies at the time of diagnosis is correlated with tumor burden but are not predictive of MCC progression after tumor surgical excision and local radiation^12,13^. In people that do not experience MCC progression, the level of T-Ag-specific antibodies declines rapidly, often becoming undetectable. In contrast, stable levels of T-Ag-specific antibodies or a brief decline followed by rapid increase usually indicates MCC progression^12–14^. These dynamics combined with the intracellular expression of T-Ag have led to a model where T-Ag-specific antibodies are produced in response to the presence of MCC but do not play a role in tumor control.

Addressing the function of tumor-specific B cells is challenging due to the diversity in tumor antigens which often varies from person-to-person even amongst cancers of the same type^15,16^. Studying tumor-specific B cell responses is further complicated by their low frequency in solid tumors^17–19^. Animal studies have demonstrated that cancer-specific B cells can enhance anti-tumor immunity by promoting antigen presentation to CD4^+^ T cells, which in turn can recruit and enhance cancer-specific CD8^+^ T cells to the tumor microenvironment^20^. Interestingly, tumors from Human Papillomavirus (HPV)^+^ head and neck cancer patients contain HPV-specific B cells ^21,22^, but whether or not these cells contribute to HPV tumor control or associate with disease outcomes is not known.

Expression of T-Ag oncoproteins by MCC allows for study of cancer-specific B cell responses across patients using T-Ag tetramers. We find that while T-Ag-specific B cells are found at increased frequencies in the blood of MCC patients compared to controls, phenotypes did not associate with disease outcome. In contrast, detection of T-Ag-specific antibody-secreting and/or germinal center B cells in tumor samples is predictive of extended progression-free survival after treatment, whereas low or undetectable levels of these cells predicted rapid MCC progression. Using antibodies and T cell receptors (TCR) identified from patient samples to engineer human T-Ag-specific B cells and CD4^+^ T cells, we demonstrate highly efficient cognate antigen presentation, suggesting a mechanism by which local specific B cells can enhance T cell-mediated immunity. Together these findings show the predictive power of local T-Ag-specific B cells as biomarkers of MCC control and suggest that functional B cell responses promote anti-tumor immunity by facilitating antigen-presentation to T cells.

## Results

### Validation of T-Ag protein tetramers to detect T-Ag-specific B cells

To identify T-Ag-specific B cells using flow cytometry, fluorescent T-Ag tetramers were used in combination with control tetramers^23,24^ to exclude B cells specific for streptavidin, the fluorochrome, and the glutathione S-transferase (GST) purification tag added to T-Ag. We used the common domain shared by small and large T-Ag isoforms since this domain is rarely mutated from patient-to-patient and most T-Ag-specific serum antibodies appear to target this region^12,25^. In contrast, the large T-Ag varies from patient-to-patient as a protein ranging from 228 - 778 amino acids depending upon the location of truncation^25–28^. Therefore, use of a tetramer only containing the “common” T-Ag domain^29,30^ allowed direct comparison of B cells binding the same antigen across cohorts. As done previously^23,24^, rare tetramer-binding B cells were more easily assessed when enriched using anti-fluorochrome microbeads before analysis. To validate the T-Ag tetramer, we immunized mice with T-Ag in complete Freund’s adjuvant (CFA) and found a robust expansion of T-Ag tetramer-binding live B cells compared to control animals injected with CFA alone (**Figure 1a, b**). Antibody-secreting cells and germinal center B cells were also present within this expanded population in T-Ag/CFA injected animals but absent in control animals (**Figure 1c**).

**Figure 1.**
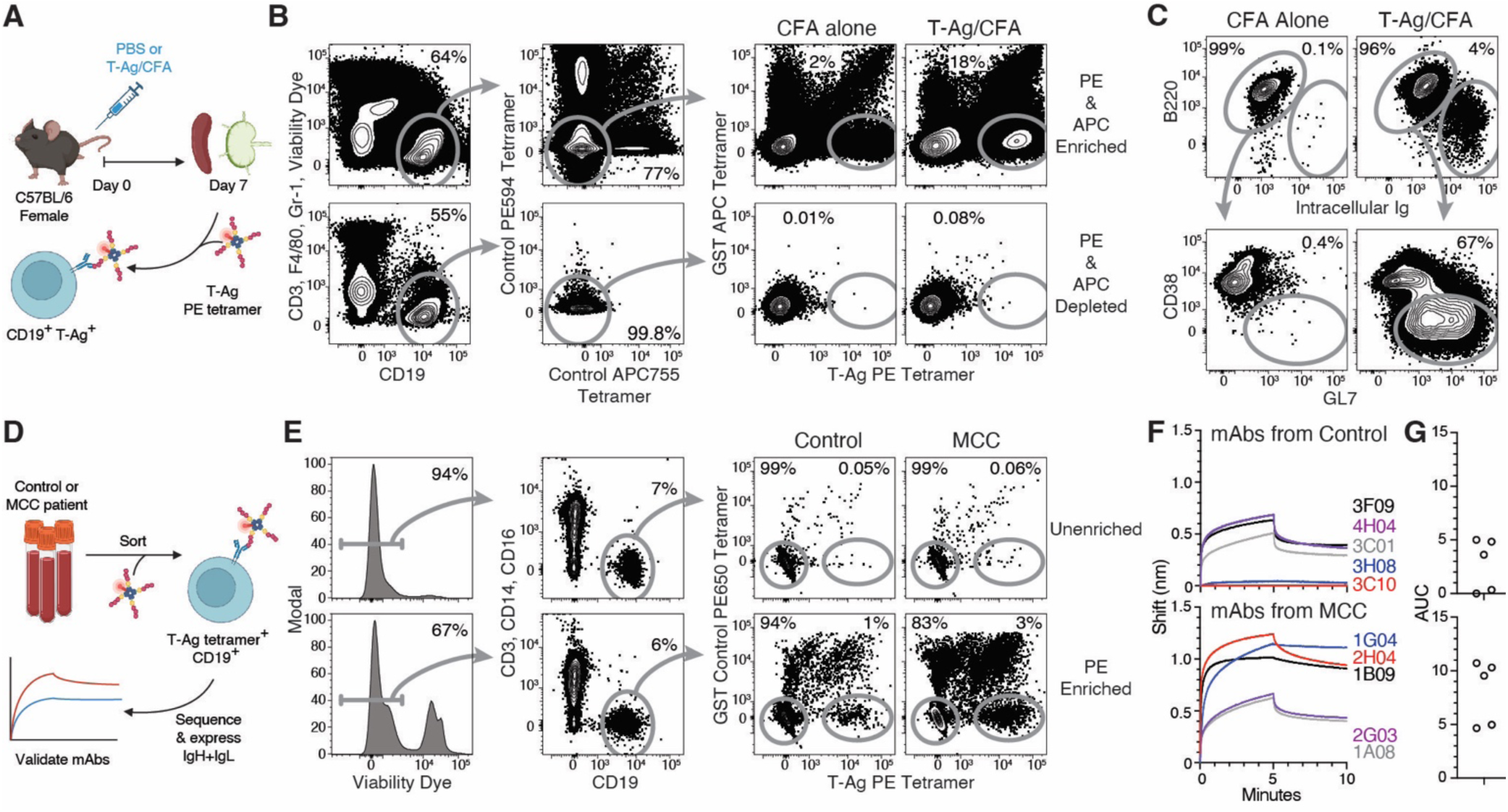
Detection of T-Ag-specific B cells in patient blood using T-Ag tetramers. **A**. Schematic representation of two experiments used to validate identification of T-Ag-specific mouse B cells using T-Ag tetramers. **B**. Representative flow cytometry of live CD19^+^ B220^+^ CD3^−^ Gr-1^−^ F4/80^−^ B cells binding T-Ag PE tetramers but not GST APC control tetramers, PE594 control tetramers or APC755 control tetramers in samples from mice seven days after subcutaneous injection of 5 μg of T-Ag in CFA in the base of the tail with compared to control mice injected with CFA alone. Each sample was enriched with anti-PE and anti-APC microbeads prior to analysis and both the PE/APC-enriched and -depleted fractions are displayed. **C**. Representative flow cytometry of T antigen tetramer-binding B220^LOW^ Ig^++^ antibody-secreting cells and GL7^+^ CD38^−^ B220^+^ Ig^+^ germinal center B cells from mice seven days after subcutaneous injection of 5 μg of T-Ag in CFA in the base of the tail compared to uninjected control mice. **D**. Schematic representation of an experiment used to validate identification of T-Ag-specific human B cells using T-Ag tetramers. **E**. Representative flow cytometry of live CD19^+^ CD3^−^ CD14^−^ CD16^−^ B cells binding T-Ag PE tetramers but not GST PE650 control tetramers in blood samples from MCC patients and controls with and without enrichment with anti-PE microbeads prior to analysis. The displayed plots contain pooled cells from three individuals with or without MCC. **F, G**. Antibodies from ten T-Ag tetramer-binding human B cells were cloned and assessed for binding to T-Ag using Bio-Layer Interferometry (BLI). Five tested antibodies derived from patient samples and five from controls. Area under the curve (AUC) of the BLI shift for each antibody. Representative of two similar experiments.

We next assessed T-Ag tetramer-binding B cells from the blood of patients with progressive T-Ag-driven MCC and control individuals with no history of MCC (**Figure 1d, e**). Single cells were sorted into individual wells of a 96-well plate and paired heavy and light chain sequences were determined using RT-PCR (**Table S1**). From these sequences five antibodies were produced from control samples and five from MCC samples. In total, 8/10 antibodies were confirmed to bind T-Ag using Bio-Layer Interferometry (BLI) indicating high specificity of the tetramer-based approach (**Figure 1f, g**).

### Frequency of antigen-experienced B cells in the blood of female patients near the time of diagnosis associates with MCC progression

After confirming that tetramer-binding B cells were T-Ag-specific, we analyzed blood cells from a prospective cohort of 47 T-Ag-driven MCC patients collected within 67 days of diagnosis and shortly before or after treatment initiation (**Figure S1**, **Table S2**). This cohort included 23 patients that experienced MCC progression within three years following our analysis, and 24 stage- and age-matched patients with non-progressive disease. In agreement with published data from larger cohorts^12,13^, the level of T-Ag-specific antibodies in blood prior to treatment was not associated with MCC progression after treatment (**Figure S2c**). Also agreeing with published data^12,13^, the level of T-Ag-specific antibodies in blood diminished in most patients that did not experience MCC progression whereas patients experiencing progressive MCC exhibited sustained or rising titers (**Figure S2d**). Combined, the alignment of blood antibody data with previous work indicates the suitability of this prospective MCC patient cohort for B cell analysis.

T-Ag-specific B cells represented ∼0.09% of B cells in blood from MCC patients, which was increased ∼10-fold compared to control samples from individuals with no history of MCC (**Figure S2e**). While the frequency of T-Ag-specific B cells ranged from 0.04 - 0.16% from patient-to-patient, there was no association with the frequency of T-Ag-specific B cells and MCC progression (**Figure S2e**). There was also no association between the frequency of T-antigen-specific B cells with the total level of T-Ag-specific antibodies in the blood (**Figure S3**).

We next considered whether different B cell subtypes associated with disease outcome and stratified the cohorts by sex because B cell phenotypes are affected by estrogen levels^31^. We first assessed total antibody-secreting cells, which varied from 0.1 – 19% of B cells in blood and frequencies did not associate with outcome (**Figures S2f, S4a**). T-Ag-specific antibody-secreting cells in blood were usually below the limit of detection, and frequencies did not associate with outcome (**Figure S4b**). In contrast, isotype-switched cells ranged from 3 - 54% of B cells and higher frequencies associated with MCC progression in female patients, but not in male patients (**Fig S2f, g**). These associations were maintained across several isotype-switched cell subtypes including CD27^+^ memory B cells and CD11c^+^ memory B cells (**Figure S2g**). The % of B cells that were isotype-switched and expressed CD71, which indicates recent activation^21^, was increased in both male and female MCC patients but did not significantly associate with progression (**Figure S2g**). Intriguingly, the frequency of T-Ag-specific B cells within each of these B cell subtypes was low and did not associate with MCC outcome, even when stratified by sex (**Figure S2h, S4b**). Together, our results demonstrate that while the level of T-Ag-specific antibodies or T-Ag-specific B cells did not associate with MCC outcome, higher frequencies of total circulating isotype-switched B cells prior to treatment associated with worse outcomes in female patients following MCC treatment.

### T-Ag-specific B cells are present in MCC tumors

The association between isotype-switched B cells in the blood with MCC progression for some patients led us to consider whether stronger associations would be identified studying B cells from tumor samples. For this we assessed a cohort of 19 patients with T-Ag-driven MCC from which single cell tumor digests were cryopreserved (**Figure 2a**). Detailed patient histories were accounted for to identify adverse clinical risk factors at the time of tumor removal that may confound survival analyses (**Figure S5**, **Table S3**). Notably, 16 of the 19 tumor samples came from male patients, limiting stratification by sex. Using known recurrence risk factors^32^, eleven patients from this cohort were considered to have low or medium risk for MCC progression, whereas eight patients were considered high risk. These risk categories we unable to predict outcome within this cohort (**Figure 2b**). As expected, T-Ag-specific antibodies at time of surgery also did not predict outcomes and were sustained or increased in patients with progressive MCC (**Figure 2c, d**). Together, these results highlight the need for additional biomarkers for more accurate MCC outcome prediction.

**Figure 2.**
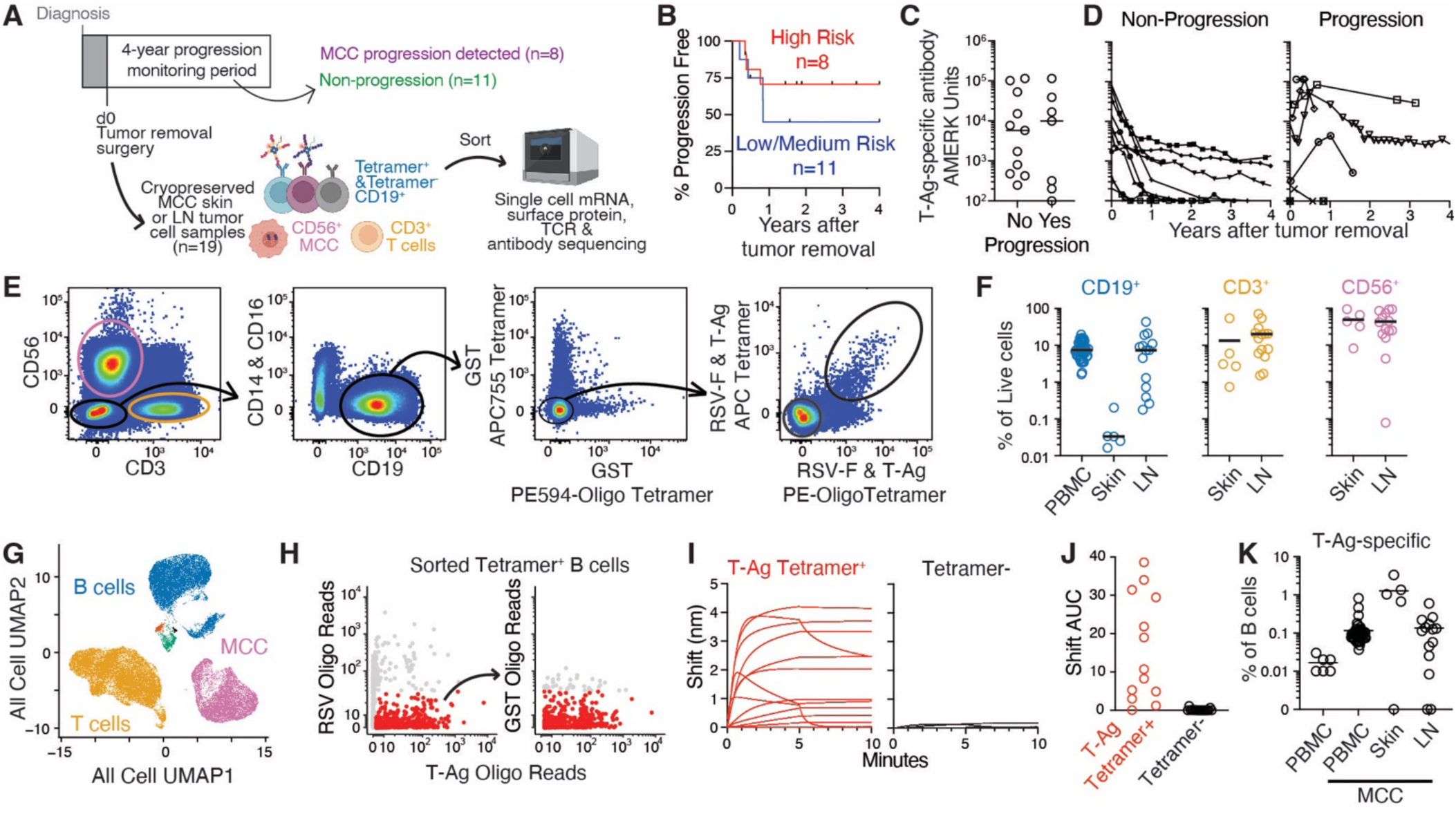
Identification and analysis of T-Ag-specific B cells from T-Ag-driven MCC patient tumor samples. **A**. Schematic representation of single cell RNA-sequencing experiments assessing B cells, T cells and MCC cells from skin tumor and LN tumor samples. **B**. Kaplan-Meier plots displaying percent progression-free survival for patients during the four-year monitoring period following tumor removal surgery stratified based on clinical risk scoring of data available at the time of surgery. **C**. Level of T-Ag-specific antibodies in the blood of MCC tumor cohort patients near the time of diagnosis stratified based upon whether patients experienced MCC progression (n=8) or not (n=11) during the four-year monitoring period. **D**. Longitudinal analysis of T-Ag-specific antibodies in the blood of MCC tumor cohort patients. Two patients from the non-progressive group for which antibody levels were not available for at least a year after diagnosis were excluded. **E**. Representative flow cytometry of CD3^+^ T cells, CD56^+^ MCC cells, and CD19^+^ B cells that bound pooled T-Ag and RSV F APC and PE-oligo tetramers but not APC755 and PE594-oligo control tetramers and B cells of unknown specificity that did not bind any tetramers. **F**. Pooled data from seven experiments showing the % of CD19^+^ B cells, % of CD3^+^ T cells, and % of CD56^+^ MCC cells in skin tumor (n=5), and LN tumor (n=14) samples from MCC patients determined by flow cytometry. **G**. UMAP clustering of CITEseq data of cells pooled from all skin and LN tumor samples after alignment and normalization with highlighted clusters of T cells (yellow), B cells (blue), MCC cells (pink), and contaminating cells (red, green). **H**. Computational strategy to identify T-antigen tetramer-binding B cells. **I, J**. Antibodies from thirteen T-Ag tetramer-binding B cells and ten from control B cells were cloned and assessed for binding to T-Ag using BLI. Representative of two similar experiments. **K**. Pooled data from seven experiments displaying the % of CD19^+^ B cells specific for T-Ag in MCC skin and LN tumor samples compared to the frequency found in blood.

To study T-Ag-specific B cell responses in MCC tumors, we performed paired Cellular Indexing of Transcriptomics and Epitopes by Sequencing (CITEseq) (**Tables S4, 5**) and antibody *IgH* + *IgL* sequencing on five MCC skin tumor samples and fourteen MCC lymph node (LN) tumor samples. Importantly, the presence of MCC in the LN samples was first detected during biopsy and later confirmed during single cell sorting based on the increased presence of CD56^+^ cells (**Figure 2e, f**). From these tumor samples we FACS-purified CD3^+^ T cells, CD56^+^ MCC cells, and CD19^+^ B cells that bound T-Ag-tetramers or Respiratory Syncytial Virus (RSV) F antigen-tetramers but not control tetramers, and B cells of unknown specificity that did not bind tetramers (**Figure 2e**). Expectedly, frequencies of CD19^+^ B cells were lower in MCC skin tumor samples when compared to blood or LN tumor samples (**Figure 2f**). In contrast, the frequency of CD3^+^ T cells and CD56^+^ MCC cells were similar across both tumor sample types (**Figure 2f**). While the frequencies of B cells, T cells, and MCC cells varied greatly from sample-to-sample, associations with MCC progression were not detected (**Figure S6a**).

After standard processing and quality control metrics, we used uniform manifold approximation and projection (UMAP)^33^ of single cell transcriptomic data from 1,015 immune-associated genes (**Table S4**), which identified clusters of MCC cells, T cells, and B cells (**Figures 2g, S7**). Within the B cell population, we identified cells that bound T-Ag tetramer-oligo but not RSV F or GST control tetramers (**Figure 2h**). We validated the specificity of our approach by confirming T-Ag binding of 87% (12/14) of antibodies derived from T-Ag tetramer-oligo^+^ B cells by BLI whereas none of the 10 antibodies cloned from tetramer^−^ B cells bound T-Ag (**Figure 2i, j, Table S6**). Together, we analyzed 941 T-Ag-specific B cells, 36,641 B cells of unknown specificity, 58,708 CD3^+^ T cells, and 33,573 CD56^+^ MCC cells from MCC tumor samples. Within the cohort, 0.01 - 4% of B cells were T-Ag-specific in the 16 of 19 samples where these cells were detected (**Figure 2k**). Despite this high variability, no association between the frequency of T-Ag-specific CD19^+^ B cells and disease progression was detected (**Figure S6b**).

### T-Ag-specific antibody-secreting cells in MCC tumor samples associates with MCC control

To assess B cell subtypes in patient tumor samples, B cells were subclustered and populations corresponding to antibody-secreting cells, germinal center B cells, memory B cells, and naïve B cells were identified (**Figure 3a, b**). B cells that did not cluster with these canonical subsets were also present and did not appear to be activated B cells since CD71 expression was not detected (**Figure 3a, b**). All analyzed B cell subtypes were observed in T-Ag-specific B cells and the much more numerous B cells of unknown specificity, with highly varied frequency distributions across the cohort (**Figure 3c**). Notably, two LN tumor samples contained a concurrent B cell malignancy (**Figure 3a, b**), Mantle Cell Lymphoma (MCL) or Chronic Lymphocytic Leukemia (CLL). For these, MCL and CLL cells were excluded to focus only on non-malignant B cells for subtype analyses.

**Figure 3.**
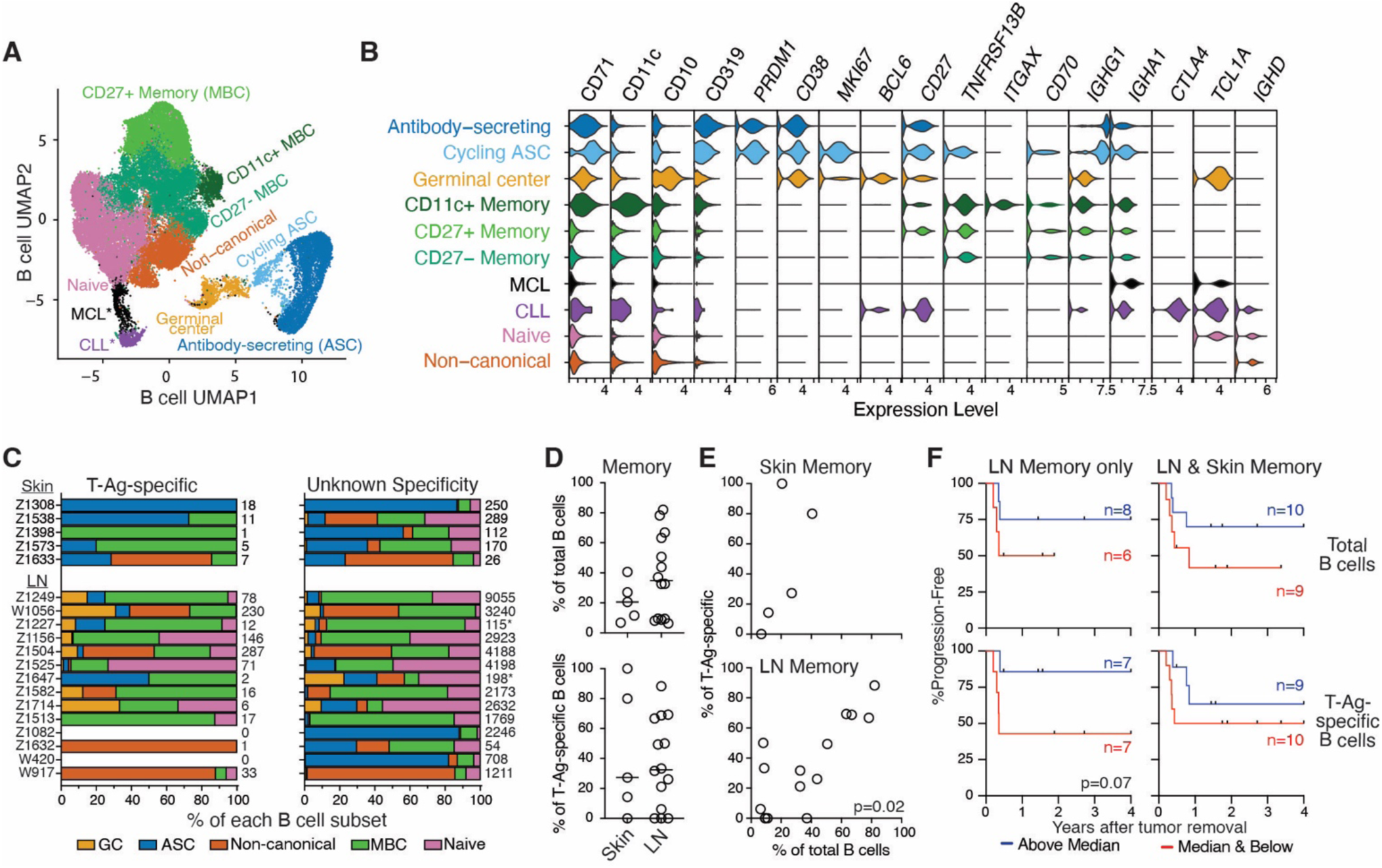
Phenotypes of T-Ag-specific B cells in MCC tumors. **A**. UMAP clustering of single cell RNA expression by B cells from pooled skin (n=5) and LN (n=14) tumor samples with clusters corresponding to different B cell subtypes highlighted with different colors. **B**. Violin plots showing expression of select genes used to identify B cell subtypes. **C**. Stacked bar graphs displaying the % of T-Ag-specific and B cells of unknown specificity of each subtype found in individual patients. The numbers to the right of each stacked bar indicate the number of cells assessed in that stacked bar. *Indicates samples in which CLL or MCL cells were excluded from subset analysis. **D**. % of total (top) and T-Ag-specific (bottom) B cells that were memory in skin and LN tumor samples. The bars indicate median for each tissue type. **E**. % of T-Ag-specific B cells that were memory versus % of total B cells that were memory in each skin (top) and lymph node (bottom) tumor samples. Correlation p value was determined using nonparametric Spearman’s test. **F**. Kaplan-Meier plots displaying % progression-free survival for patients during the monitoring period following analysis divided into groups with a % of total (top) and T-Ag-specific (bottom) B cells that were memory above (blue) or at/below (red) the median. The p value nearing significance generated using Mantel-Cox Log-rank test.

Since increased frequencies of isotype-switched B cells in the blood associated with MCC progression for some patients, we next assessed memory B cells in tumor samples as a population able to exit the tissue and enter the blood. Memory B cells were detected in every tumor sample and ranged from 6 - 82% of B cells (**Figure 3d**). T-Ag-specific memory B cells exhibited similar variability but were not detected in three LN tumor samples and one skin tumor sample (**Figure 3d**). In LN tumor samples, the frequency of T-Ag-specific memory B cells correlated with the total frequency of memory B cells (**Figure 3e**). Higher frequency of T-Ag-specific or total memory B cells in tumor samples did not associate with MCC progression (**Figure 3f**). Since MCC skin lesion samples were limited, these analyses were conducted with LN tumor samples alone or combined with skin tumor samples (**Figure 3f**). Intriguingly, lower frequency of T-Ag-specific memory B cells in LN tumor samples trended towards an association with MCC progression but did not achieve statistical significance (**Figure 3f**).

Antibody-secreting cells also exhibited high sample-to-sample variability, accounting for 10 - 87% and 1 - 85% of CD19^+^ cells in skin and LN tumor samples, respectively (**Figure 4a**). Variability was similar within the T-Ag-specific B cell population, except that T-Ag-specific antibody-secreting cells were not detected in one skin tumor sample and half of the fourteen LN tumor samples (**Figure 4a**). Unlike memory B cells (**Figure 3e**), higher frequency of T-Ag-specific antibody-secreting cells did not correlate with frequencies of total antibody-secreting cells in MCC tumor samples (**Figure 4b**).

**Figure 4.**
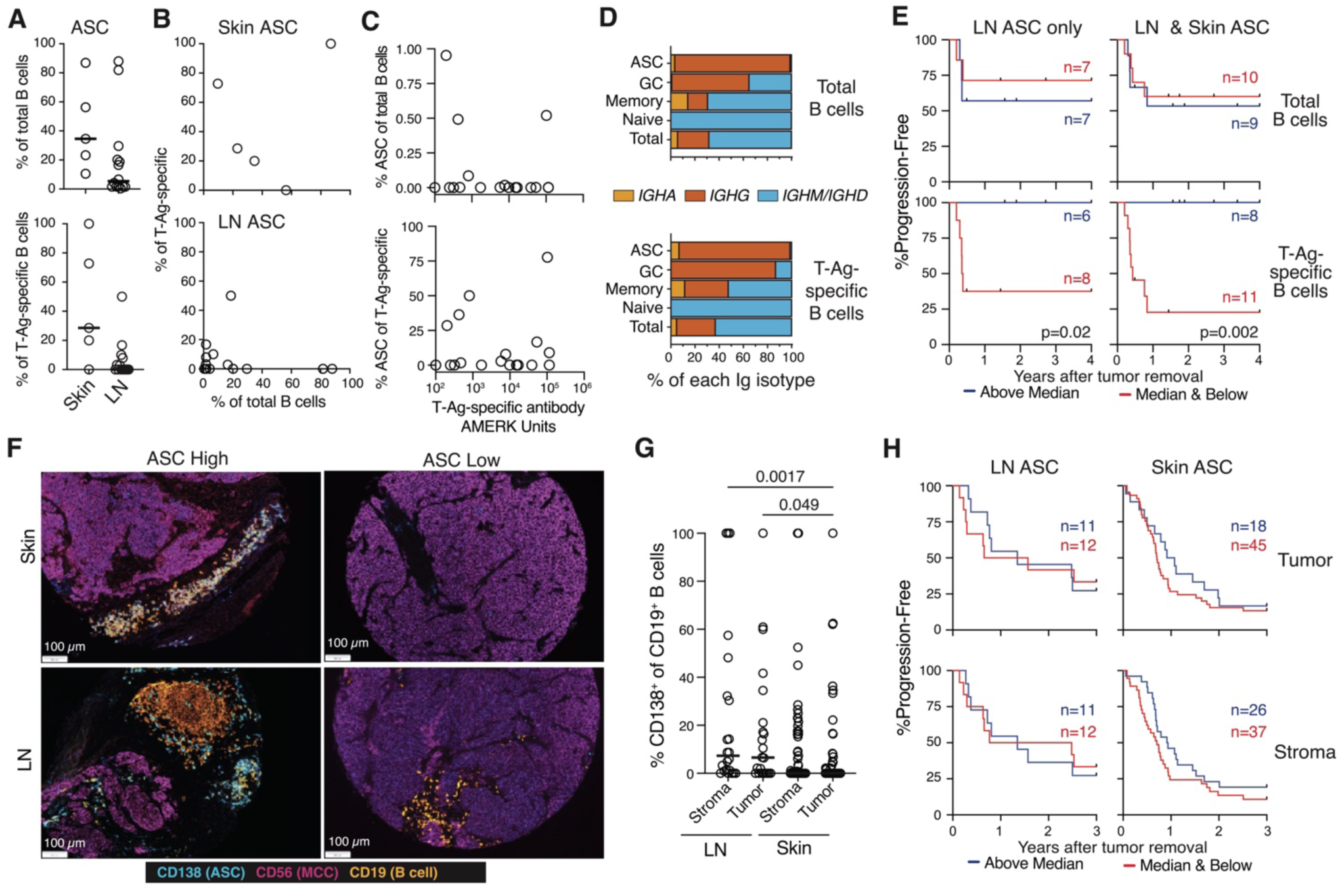
Lower frequency of T-Ag-specific antibody-secreting cells in MCC tumors associates with progressive disease. **A**. Data from single cell sequencing experiments showing the % of total (top) and T-Ag-specific (bottom) antibody-secreting cells in skin and LN tumor samples. Bars indicate medians. **B**. % of T-Ag-specific cells that were antibody-secreting versus the % of all B cells that were antibody-secreting in skin (top) and LN (bottom) tumor samples. **C**. Level of T-Ag-specific antibodies in the blood of MCC patients near the time of surgery versus the % of T-Ag-specific or total B cells that were antibody-secreting in skin tumor (top) and LN (bottom) tumor samples. **D**. % of B cells of total (top) or T-Ag-specific B cells of the listed B cell subtype that expressed *IGHM/IGHD* or class-switched to *IGHA* or an *IGHG.* **E**. Kaplan-Meier plots displaying % progression-free survival for patients during the monitoring period divided into groups with a % of total (top) and T-Ag-specific (bottom) B cells that were antibody-secreting above (blue) or at/below (red) the median. Significant p values generated using Mantel-Cox Log-rank test are displayed. **F**. Select mIHC images of skin and lymph node tumor samples determined to contain high or low frequencies of CD19^+^ CD138^+^ antibody-secreting cells. **G**. % of CD19^+^ cells that were CD138^+^ antibody-secreting cells within the CD56^+^ tumor tissue or adjacent non-tumor stroma detected using mIHC of LN (n=23) and skin tumor samples (n=63). **H**. Kaplan-Meier plots displaying % progression-free survival for patients during a three-year monitoring period following analysis stratified based on % of B cells that were antibody-secreting above (blue) or at/below (red) the median within the tumor tissue (top) and adjacent non-tumor stroma (bottom) determined by mIHC.

Similarly, the frequency of T-Ag-specific antibody-secreting cells in MCC tumor samples did not correlate with levels of T-Ag-specific antibodies in the blood at the time of tumor removal (**Figure 4c**). The lack of correlation between levels of antibody in blood and frequency of detected antibody-secreting cells did not appear to be the result of IgG being detected in blood and antibody-secreting cells of all isotypes being interrogated since nearly all antibody-secreting cells detected were IgG^+^ (**Figure 4d**). Combined, these data suggested that the antibody-secreting cell response detected in tumor samples is distinct from what can be inferred from analysis of serum antibodies in the blood.

We next considered whether antibody-secreting cell frequencies in tumor samples associated with MCC outcome after treatment. Since MCC skin lesion samples were limited, we conducted separate analyses either with LN tumor samples alone or combined with skin tumor samples. Strikingly, we found the presence of detectable T-Ag-specific antibody-secreting cells in LN tumor samples was associated with extended progression-free survival following tumor removal (**Figure 4e**). Similar results were found when skin tumor samples were included based upon whether the frequency of T-Ag-specific antibody-secreting cells was above or below the median found in skin tumors (**Figure 4e**). In contrast, a higher frequency of total antibody-secreting cells in skin or LN tumors was not associated with MCC outcome (**Figure 4e**). To further probe relationships between antibody-secreting cells and MCC progression, we used multiplex immunohistochemistry (mIHC) to assess an independent cohort of 23 MCC LN tumor and 63 MCC skin tumor sections (**Figure 4f, S8**, **S9, Table S7**). As observed by CITEseq, the frequency of CD138^+^ antibody-secreting cells within CD19^+^ cells in MCC tumors varied widely but did not differ between regions of tumor mass and adjacent non-tumor tissue stroma (**Figure 4f, g**). Similarly, the frequency of antibody-secreting cells out of total B cells tumor mass or adjacent non-tumor tissue stroma did not associate with MCC outcomes for skin or LN tumor samples (**Figure 4h**). Together, our results indicate that while high frequencies of antibody-secreting cells in MCC tumor samples does not associate with MCC outcome, the absence of detectable T-Ag-specific cells within this population associates with MCC progression.

### Germinal center response in MCC tumor samples associates with MCC control

The presence of tertiary lymphoid structures able to support germinal centers in tumors associates with better outcomes for many solid cancers^17–19^. We identified T-Ag-specific germinal center B cells in eight of the fourteen LN tumor samples but failed to detect any in skin tumor samples (**Figure 5a**). Few germinal center B cells were detected in skin tumor samples overall (**Figure 5a**) in agreement with previous work demonstrating limited development of organized tertiary lymphoid structures in MCC skin tumor samples^34–36^. Among the LN tumor samples, higher frequencies of T-Ag-specific germinal center B cells trended with higher frequencies of total germinal center B cells (**Figure 5b**) even though T-Ag-specific cells made up less than 1% of germinal center B cells when detected (**Figure 5c**). Remarkably, the presence of detected T-Ag-specific germinal center B cells in LN tumor samples was predictive of prolonged progression-free survival, whereas absence of these cells associated with rapid disease-progression (**Figure 5d**). Interestingly, higher frequencies of total germinal center B cells in LN tumor samples also associated with progression-free survival (**Figure 5d**).

**Figure 5:**
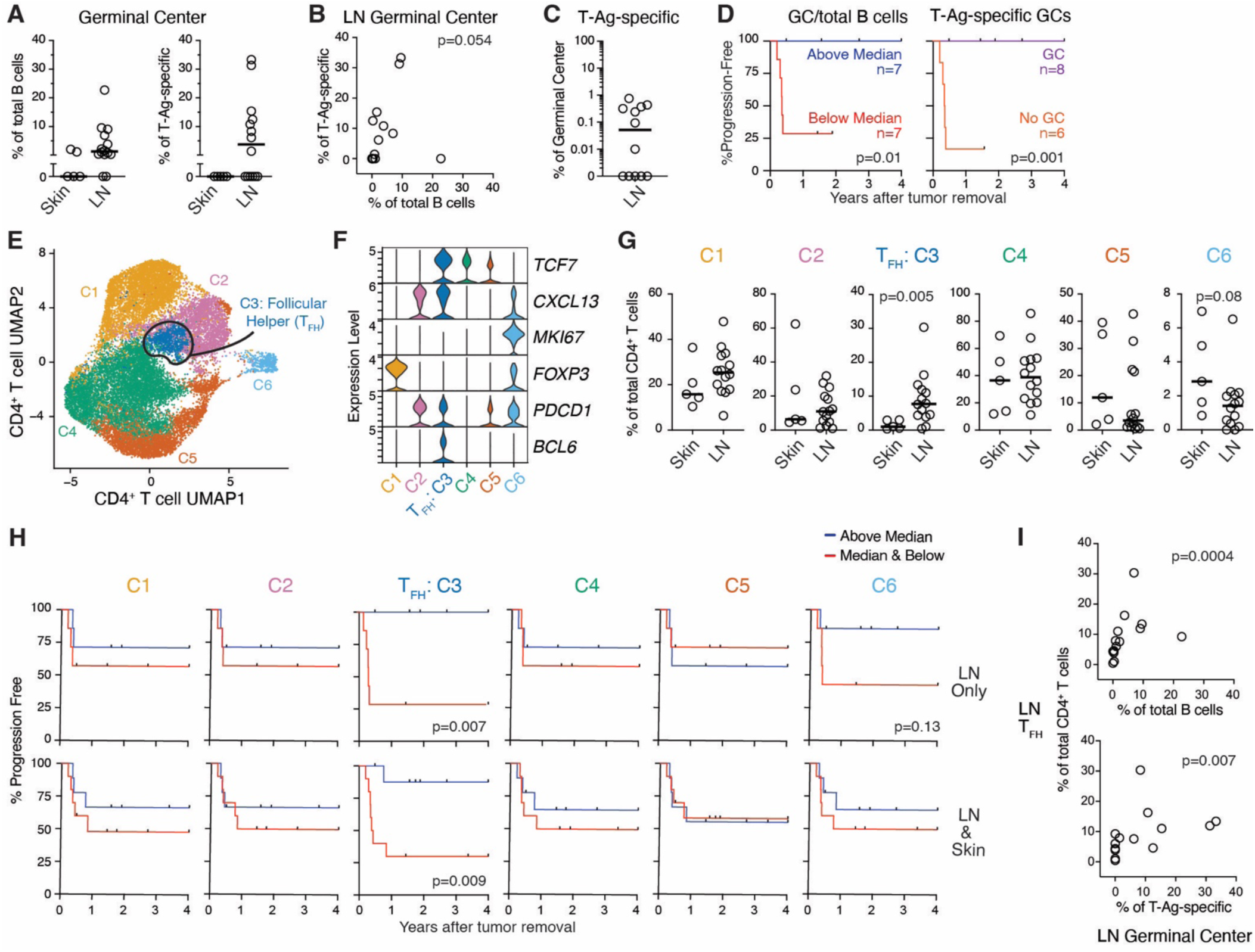
Lower frequency of germinal center B cells and T_FH_ cells in MCC tumor samples associates with progressive disease. **A**. Data from single cell sequencing experiments showing the % of total (top) and T-Ag-specific (bottom) B cells that were germinal center phenotype in skin and LN tumor samples. Bars indicate medians. **B**. % of T-Ag-specific B cells that were germinal center phenotype versus the % of all B cells that were germinal center phenotype in LN tumor samples. Correlation p value determined using nonparametric Spearman’s test. **C**. % of T-Ag-specific cells within the total germinal center B cell population in LN tumor samples. **D**. Kaplan-Meier plots displaying % progression-free survival for patients during the four-year monitoring period following analysis divided into groups with a % of total B cells (left) that were germinal center phenotype above (blue) or at/below (red) the median and whether T-Ag-specific (right) germinal center phenotype cells were detected (violet) or below the limit of detection (orange). Displayed p values generated using Mantel-Cox Log-rank test. **E**. UMAP clustering of single cell RNA expression by CD4^+^ T cells from pooled skin (n=5) and LN (n=14) tumor samples with clusters corresponding to subtypes highlighted with different colors. **F**. Violin plots showing the expression of select genes used to identify T_FH_ cells. **G**. % of CD4^+^ T cells that were in each cluster within skin and LN tumor samples. Bars indicate median and significant p value generated using a non-parametric Mann-Whitney test is displayed. **H**. Kaplan-Meier plots displaying % progression-free survival for patients during the four-year monitoring period following analysis divided into groups with a % of CD4^+^ T cells that were in each cluster above (blue) or at/below (red) the median. Significant p values generated using Mantel-Cox Log-rank test are displayed. **I**. % of total (left) and T-Ag-specific (right) B cells that were germinal center phenotype versus the % of CD4+ T cells that were T_FH_ in LN tumor samples. Correlation p values determined using nonparametric Spearman’s test.

Because robust germinal center B cell responses are dependent on signals from follicular helper CD4^+^ T cells (T_FH_), we assessed whether this T cell population associated with MCC outcome. Sub-clustering of CD4^+^ T cells revealed a population of cells with a T_FH_ signature^37^ that included increased expression of *BCL6*, *PDCD1*, *CXCL13*, and *TCF7* transcripts (**Figure 5e, f**). CD4^+^ T cells in this cluster were detected in both skin and LN tumor samples but were more abundant in LN (**Figure 5g**). Like the germinal center B cell response, frequencies of T_FH_ cells above the median in LN tumor samples were also predictive of progression-free survival, whereas patients whose tumors had less T_FH_ cells were more likely to have rapid MCC progression (**Figure 5h**). Expectedly, tumor samples with more T_FH_ cells tended to have higher frequencies of T-Ag-specific and total germinal center B cells (**Figure 5i**). In contrast, none of the five other CD4^+^ T cell clusters associated with disease outcome (**Figure 5g, h**).

### T-Ag-specific B cells potently activate T-Ag-specific CD4^+^ T cells

To confirm the presence of T-Ag-specific T_FH_ cells in MCC tumors, we examined paired TCRα/β CDR3 sequences from CD4^+^ T cells from three patient tumors. This analysis revealed that 31% (408/1315) of these cells shared identical paired TCRα/β CDR3 sequences with at least one other cell, indicating clonal expansion (**Figure 6a, Table S8**). TCRs from 69 clonal families containing *CXCL13^+^* members were expressed in primary CD4^+^ T cells derived from healthy donors using a previous described lentiviral transduction approach^37^ (**Figure 6b**). Transduced T cells were next assessed for T-Ag specificity by incubation with autologous B cells in the presence and absence of T-Ag peptides. This assay revealed that 23% (16/69) of tested clonal TCRs secreted IFNγ in response to pools of T-Ag peptides (**Figure 6c, S10, Table S9**), confirming specificity for T-Ag.

**Figure 6.**
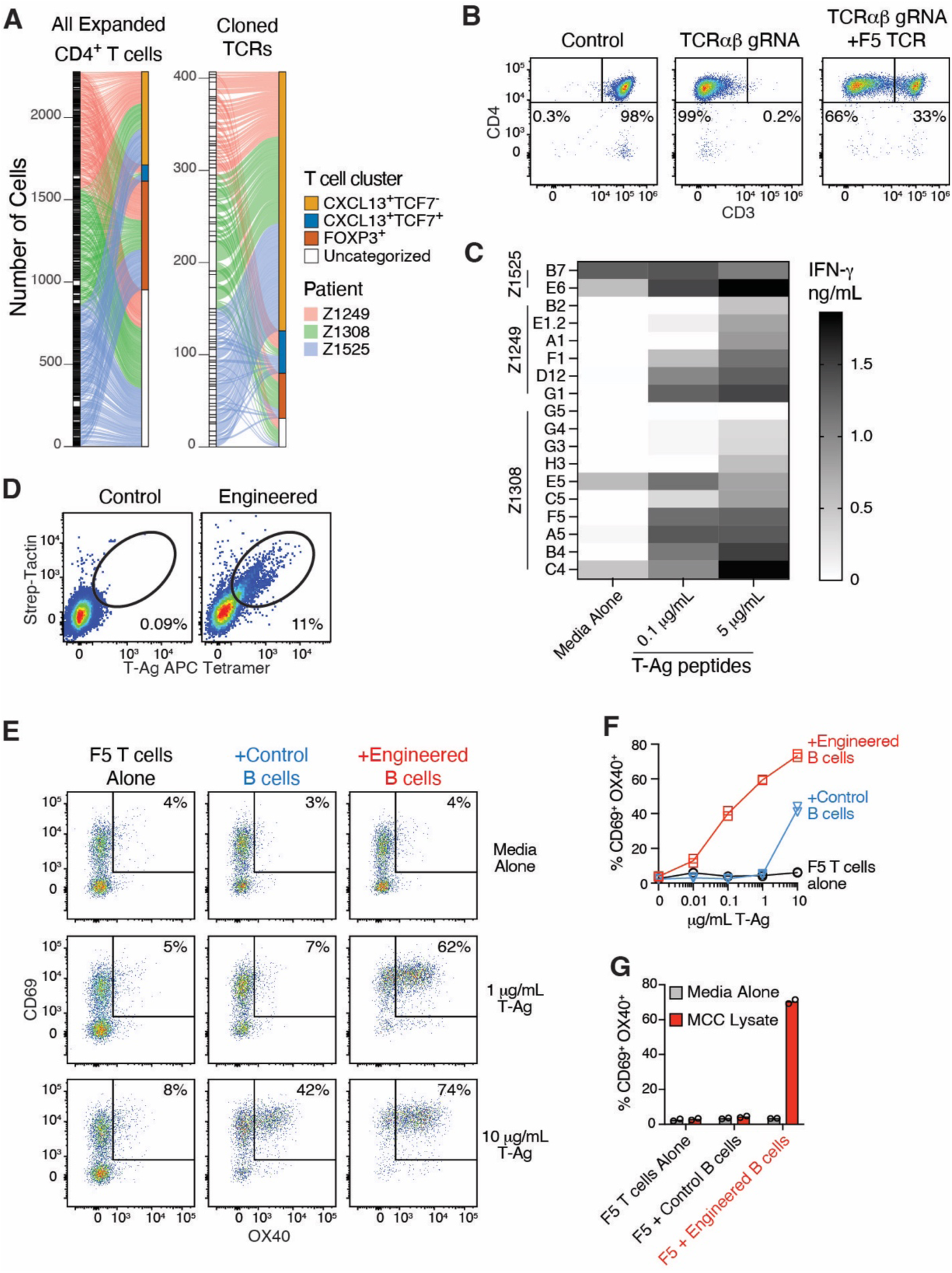
T-Ag-specific engineered B cells potently stimulate T-Ag-specific engineered CD4^+^ T cells. **A**. TCRs expressed by all CD4^+^ T cells (left) and identical TCRs expressed by multiple cells (right) from three LN and skin tumor samples linked to cell subtype. **B**. Representative flow cytometry of the expression of lentiviral-expressed T-Ag-specific TCR in CD4^+^ from blood detected as the expression of CD3 on the cell surface in cells lacking endogenous TCR due to CRISPR knockout using gRNA targeting TCRα and TCRβ. **C**. Eighteen clonal T_FH_ TCRs from MCC tumor samples were expressed via lentivirus in CD4^+^ cells from blood and incubated with sample-matched B cells in the absence or presence of 0.1 or 5 μg/mL of T-Ag peptide pools for three days prior to assessment of supernatant for secreted IFNγ. Representative of two similar experiments. **D**. Representative flow cytometry of Strep-Tactin and T-Ag binding to human B cells engineered to express a T-Ag-specific antibodies using CRISPR/Cas9. Antibody constructs encoding 1G04, 2H04, and 1B09 from containing the full antibody light chains physically linked to the heavy chain VDJs with a linker^39^ containing Strep-tagII^40–42^ were pooled prior to engineering. **E**. Representative flow cytometry and (**F**) quantitation of CD69 and OX40 expression by lentiviral-transduced T-Ag-specific F5 TCR CD4^+^ T cells expressing following 72-hour co-culture with CRISPR-engineered T-Ag-specific oligoclonal B cells or control B cells in the presence or absence of various concentrations of T-Ag. **G**. Quantitation of CD69 and OX40 expression by lentiviral-transduced T-Ag-specific F5 TCR CD4^+^ T cells expressing the were co-cultured for 24 hours with CRISPR-engineered T-Ag-specific oligoclonal B cells or control B cells in the presence or absence of MCC cell line lysate. **E**-**G** representative of two similar experiments.

We next considered whether T-Ag-specific B cells could more potently stimulate T-Ag-specific CD4^+^ T cells compared to B cells of unknown specificity. For this, we cultured lentiviral-transduced T-Ag-specific CD4^+^ T cells with CRISPR-engineered T-Ag-specific B cells generated using a previously described approach^38^. For these assays we used a cocktail of constructs encoding the three T-Ag-specific antibodies, 1G04, 2H04, and 1B09 (**Figure S1F**) to simulate an oligoclonal B cell population. Antibody constructs containing DNA producing the full antibody light chain physically linked to the heavy chain VDJ with a 57 amino acid glycine-serine linker^39^ were pooled prior to engineering was expression was validated based upon binding T-Ag tetramer and Strep-Tactin (**Figure 6d**), which binds Strep-tagII^40–42^ included in the linker. These assays revealed that T-Ag-specific engineered B cells from a healthy donor that expressed the HLA DR4-DQ8 haplotype that presents the T-Ag could induce similar levels of CD69 and OX40 by CD4^+^ T cells expressing the T-Ag-specific TCR F5 in response to 100-fold lower concentrations of purified T-Ag compared to control B cells (**Figure 6e, f**). T-Ag-specific engineered B cells could also stimulate CD69 and OX40 by T-Ag-specific F5 TCR CD4^+^ T cells in response to MCC cell lysate from a tumor cell line, which was not found with control B cells (**Figure 6g**). Combined, these data indicate that T-Ag-specific B cells can potently activate T-Ag-specific CD4^+^ T cells.

### Antibody-secreting cells and memory B cells in tumor samples are largely germinal center-derived

Our data demonstrated that the T-Ag-specific germinal center and antibody-secreting B cell responses in MCC tumors associate with better progression-free survival (**Figure 4e, 5d**). Since antibody-secreting cell differentiation can occur through germinal-center dependent and independent pathways, we analyzed paired antibody heavy and light chain sequences and clonal dynamics of B cells in MCC tumor samples for evidence of somatic hypermutation in a germinal center.

To assess somatic hypermutation in B cells from MCC tumors, we determined the mutation frequency in *IgH* sequences with respect to germline encoded sequences. Most T-Ag-specific antibody-secreting cells and germinal center B cells had *IgH* mutations at similar levels to B cells of unknown specificity with these phenotypes (**Figure 7a**). Somatic hypermutation was also present in T-Ag-specific memory B cells found in tumor samples, even in samples where germinal center B cells were not detected (**Figure 7a**). Similarly, *IgH* mutation frequencies were similar in B cells from skin and LN tumor samples, independently of T-Ag specificity (**Figure 7b**). Notably, the frequency of somatic hypermutation did not associate with MCC progression (**Figure 7c**).

**Figure 7:**
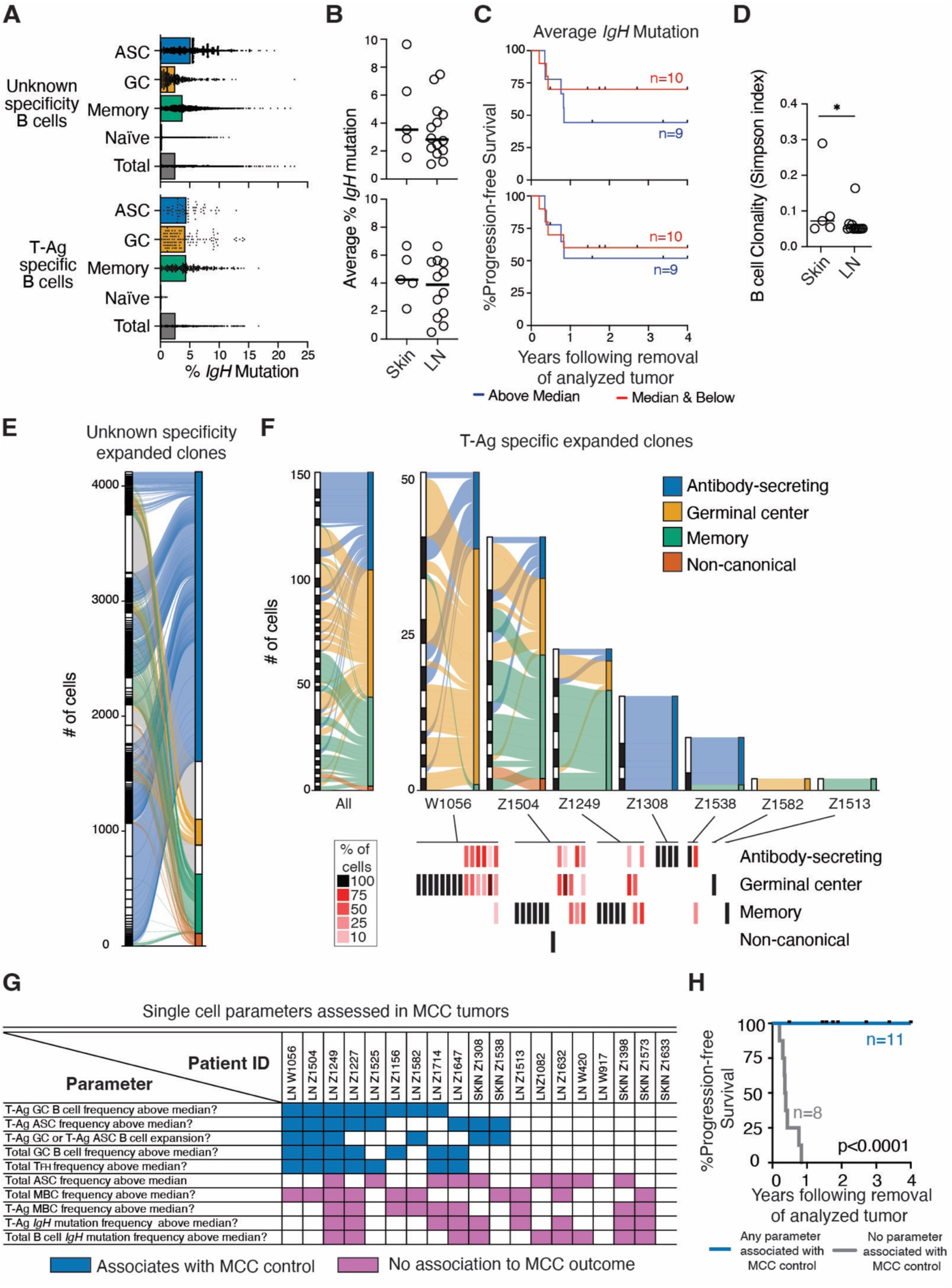
Analysis of somatic hypermutation and clonality indicates many antibody-secreting cells and memory B cells in tumors are germinal center-derived. **A**. % of mutated *IgH* nucleotides for cells within the listed subpopulations of B cells of unknown specificities (top) and T-Ag-specific B cells (bottom). **B**. Average % of mutated *IgH* nucleotides for B cells of unknown specificities (top) and T-Ag-specific B cells (bottom) found in skin and LN tumor samples. **C**. Kaplan-Meier plots displaying % progression-free survival for patients during the monitoring period following analysis divided into groups with a % mutation rate above (blue) or at/below (red) the median. **D**. Assessment of clonality within the B cells found in skin and LN tumor samples using Simpson Diversity Index. **E**. Stacked clonal lineage alluvial plot with lines linking individual cells within clonal families (left) with the B cell subtype (right) of that cell within B cells of unknown specificities. Each clonal family is displayed as an individual bar stacked in alternating black and white sized corresponding to the number of cells detected in that clonal family. **F**. Stacked clonal lineage plots (top) and heatmaps (bottom) of all T-Ag-specific clonal families from LN and skin tumor samples, and plots divided into individual patients. The columns in the heat maps correspond to individual clonal families. **G**. Summary of findings from analyses of B cells and T cells with phenotypes with significant associations with MCC control highlighted in blue. **H**. Kaplan-Meier plots displaying % progression-free survival for patients during the monitoring period following analysis divided into a group with any of the B or T cell characteristics associated with MCC control from **G** (blue), versus the group that did not have any of those characteristics. The p value generated using Mantel-Cox Log-rank test.

While somatic hypermutation strongly suggested that T-Ag-specific antibody-secreting cells and memory B cells derived from germinal centers, we sought to confirm this by analysis of clonal families containing germinal center and non-germinal center members. Analysis of paired *IgH* + *IgL* sequences revealed that B cells from skin tumor samples were slightly more clonally related than B cells from LN tumor samples (**Figure 7d**). In total, we identified 4,119 expanded B cells of unknown specificity amongst 589 clonal families spread across 17 MCC patient tumor samples (**Figure 7e**). T-Ag-specific B cell clonal families were only found in 7 of the 19 tumor samples and amounted to 150 expanded cells amongst 42 clonal families (**Figure 7f, Table S10**). Intriguingly, six of the seven samples with detectable clonal families were from patients who did not progress, with the smallest clonal family being derived from lymph node tumor sample from patient Z1513, who experienced MCC progression (**Figure 7f, S5**).

Not surprisingly, the three LN tumor samples with the most abundant T-Ag-specific B cell clonal families contained mutated antibody-secreting cells and germinal center B cells within the same clonal family (**Figure 7f**). In fact, 10 of the 18 T-antigen-specific clonal families containing an antibody-secreting B cell also contained a germinal center B cell (**Figure 7f**). These results demonstrated that T-Ag-specific antibody-secreting cells and memory B cells can derive from germinal centers found in LN tumor samples.

As a final analysis we compiled the analyses of B cells and T cells to determine if combinations of parameters better predicted disease outcome (**Figure 7g**). When assessed together, we found that tumor samples from all eight MCC patients in the cohort that that later experienced MCC progression contained low frequencies of T-Ag-specific germinal center B cells, low frequencies of T-Ag-specific antibody-secreting cells, undetectable T-Ag-specific B cell clonal families, low frequencies of total germinal center B cells, and low frequencies of T_FH_ cells (**Figure 7g, h**). In contrast, all eleven MCC patients had higher frequencies of at least two of these five B cell and T cell readouts (**Figure 7g, h**).

## Discussion

Increasing evidence suggests an important role for B cells in immune control of tumors. Here we directly analyzed B cells specific for T-Ag oncoprotein during MCC in search of biomarkers able to predict MCC progression. Similar to analyses of T-Ag-specific antibodies in the blood^12,13^, analysis of T-Ag-specific B cells in the blood near the time of diagnosis did not yield associations with disease progression. Intriguingly, increased frequencies of isotype-switched total B cells in the blood of female patients did associate with disease progression. It is intriguing to speculate that these links are related to the role estrogen plays in mediating increased levels of isotype-switched B cells in the blood^31^.

We also found increasedisotype-switched CD71^+^ B cells in the blood of both male and female MCC patients. This is not surprising given that CD71 is a marker of B cells recently activated by antigen in response to cancer and vaccination^21^. It is unclear what antigen these activated CD71^+^ B cells are recognizing since the vast majority did not bind the portion of T-Ag usedhere. Some of these activated CD71^+^ B cells could be specific for regions of large T-Ag that are not conserved from patient-to-patient. Alternatively, these activated CD71^+^ B cells could be specific for self-antigens since self-antigen-specific antibodies are found in many cancers^43,44^.

Findings from B cell populations in the blood were intriguing but not robust enough to be clinically actionable biomarkers, prompting assessment of B cells from 19 MCC patient tumor samples which revealed several predictive biomarkers of MCC progression. We found that ongoing germinal center B cell responses and high frequencies of T-Ag-specific antibody-secreting cells in tumor samples predicted control of MCC after treatment with remarkable accuracy. Given the strong predictive ability of T-Ag-specific B cell phenotypes in tumors, it was surprising that the level of T-Ag-specific antibodies in blood at the time of treatment is not predictive of outcome. We hypothesize that this difference is reflective of T-Ag-specific B cell responses we are unable to measure, such as responses in the weeks prior to tumor surgical excision, or responses in anatomical locations we are unable to assess, such as sites of disease spread or unaffected draining lymph nodes. This hypothesis is supported by our data indicating the presence of somatic hypermutation in most T-Ag-specific B cells in tumors even when germinal center responses are not detected.

Studies in other cancers analyzing the presence of antibody-secreting cells in tumors have often reported positive, negative, and no associations with disease outcome^45^. Our work studying the overall antibody-secreting cell responses using flow cytometry and immunofluorescence did not find an association with MCC progression. However, by studying T-Ag-specific B cells, we found a strong association between low frequencies of T-Ag-specific antibody-secreting cells and MCC progression. These data raise questions about the specificities and differentiation pathways of the more numerous antibody-secreting cells that are not T-Ag specific.

While we found several promising biomarkers of progressive MCC, more work is needed to understand the mechanism. Enhanced T-Ag-specific germinal center B cell and antibody-secreting cell responses in MCC tumor samples may reflect of an overall more robust immune response resulting in more tumor killing. In this scenario B cells are responding to the presence of more T-Ag released from killed MCC cells. However, if more cell-free T-Ag is present, the levels of T-Ag-specific antibodies in blood may also be expected to be increased, which is not the case. The association of T_FH_ cell frequencies with MCC control, but not other CD4^+^ T cell subsets, also suggests a role for B cells. If increased B cell responses are merely of reflection of an overall better tumor-killing immune response, a specific increase in T_FH_ cells would not be required as numerous CD4^+^ populations would be expected to be increased if merely a reflection of overall immune potency.

We hypothesize that T-Ag-specific B cells help enhance the response of T-Ag-specific T cell killing of tumors. In support of this, we show that B cells engineered to be T-Ag-specific can potently activate CD4^+^ T cells expressing a T-Ag-specific TCR derived from a patient tumor sample. While it is unlikely that CD4^+^ T cells directly kill MCC due to the lack of MHC class II expression by MCC, CD4^+^ T cells can enhance antitumor CD8^+^ T cell responses^20,46,47^. Likewise, data from a mouse model has suggested that B cell stimulation of T_FH_ cells enhances IL-21 production which in turn enhances tumor killing by CD8^+^ T cells^20^. Newly generated mouse models of MCC^48^ will likely help to answer these questions.

## Abbreviations

APC: allophycocyanin
APC755: APC-DyLight 755
ASC: antibody-secreting cell
AUC: area under the curve
BLI: Bio-Layer Interferometry
CFA: complete Freund’s adjuvant
CITEseq: Cellular Indexing of Transcriptomics and Epitopes by Sequencing
CLL: Chronic Lymphocytic Leukemia
GST: glutathione S-transferase
HPV: Human Papillomavirus
Ig: immunoglobulin
*IgH*: antibody heavy chain gene
*IgL*: antibody light chain gene
LN: lymph node
MCC: Merkel cell carcinoma
MCL: Mantle Cell Lymphoma
MCPyV: Merkel cell polyomavirus
mIHC: multiplex immunohistochemistry
PBMC: peripheral blood mononuclear cells
PE: R-phycoerythrin
PE594: PE-DyLight 594
PE650: PE-DyLight 650
RSV: Respiratory Syncytial Virus
T-Ag: T-antigen
TCR: T cell receptor
T_FH_: follicular helper CD4^+^ T cell
UMAP: uniform manifold approximation and projection.

---

**Supplemental Figure 1.**
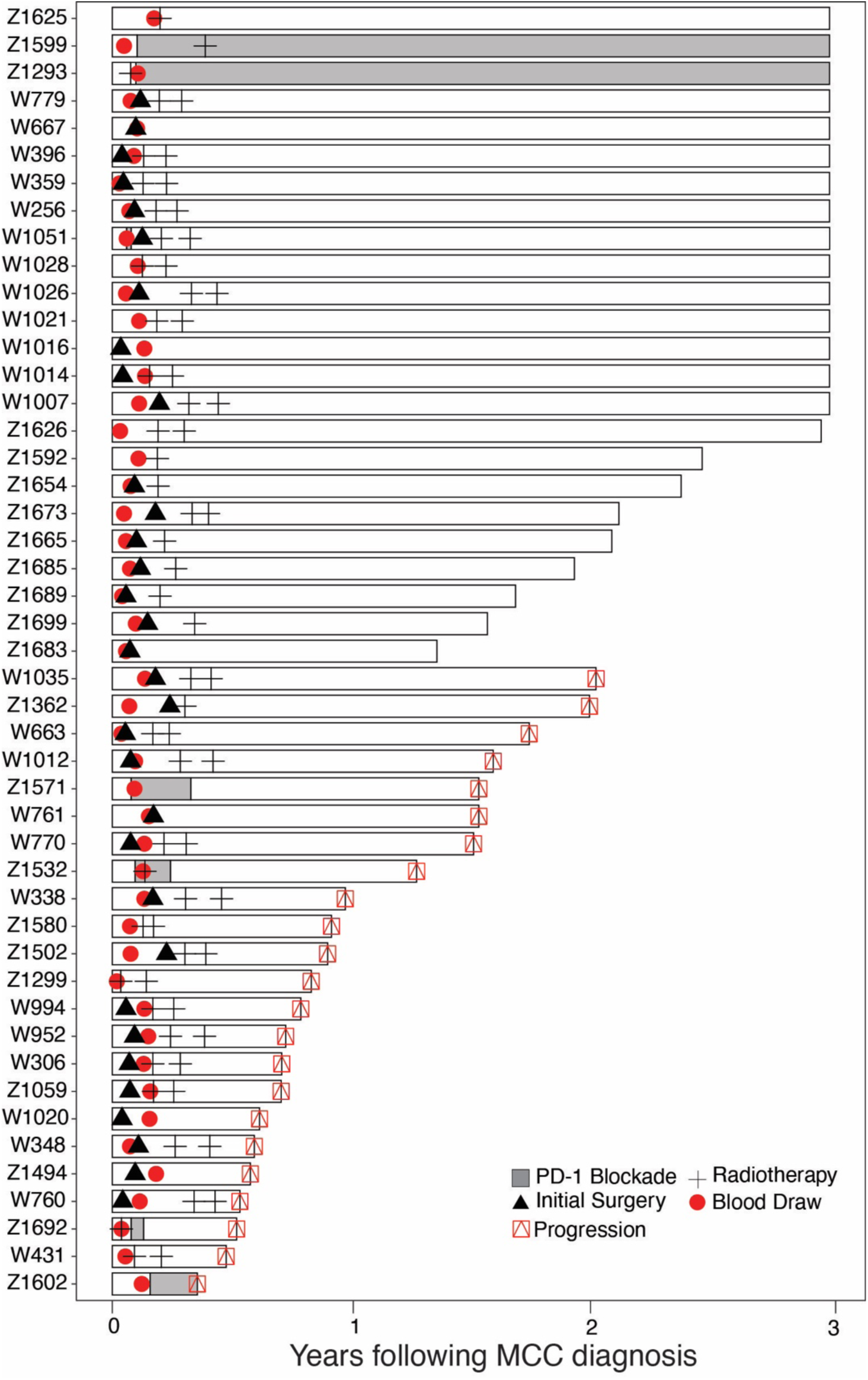
PBMC cohort patient history summaries.

**Supplemental Figure 2.**
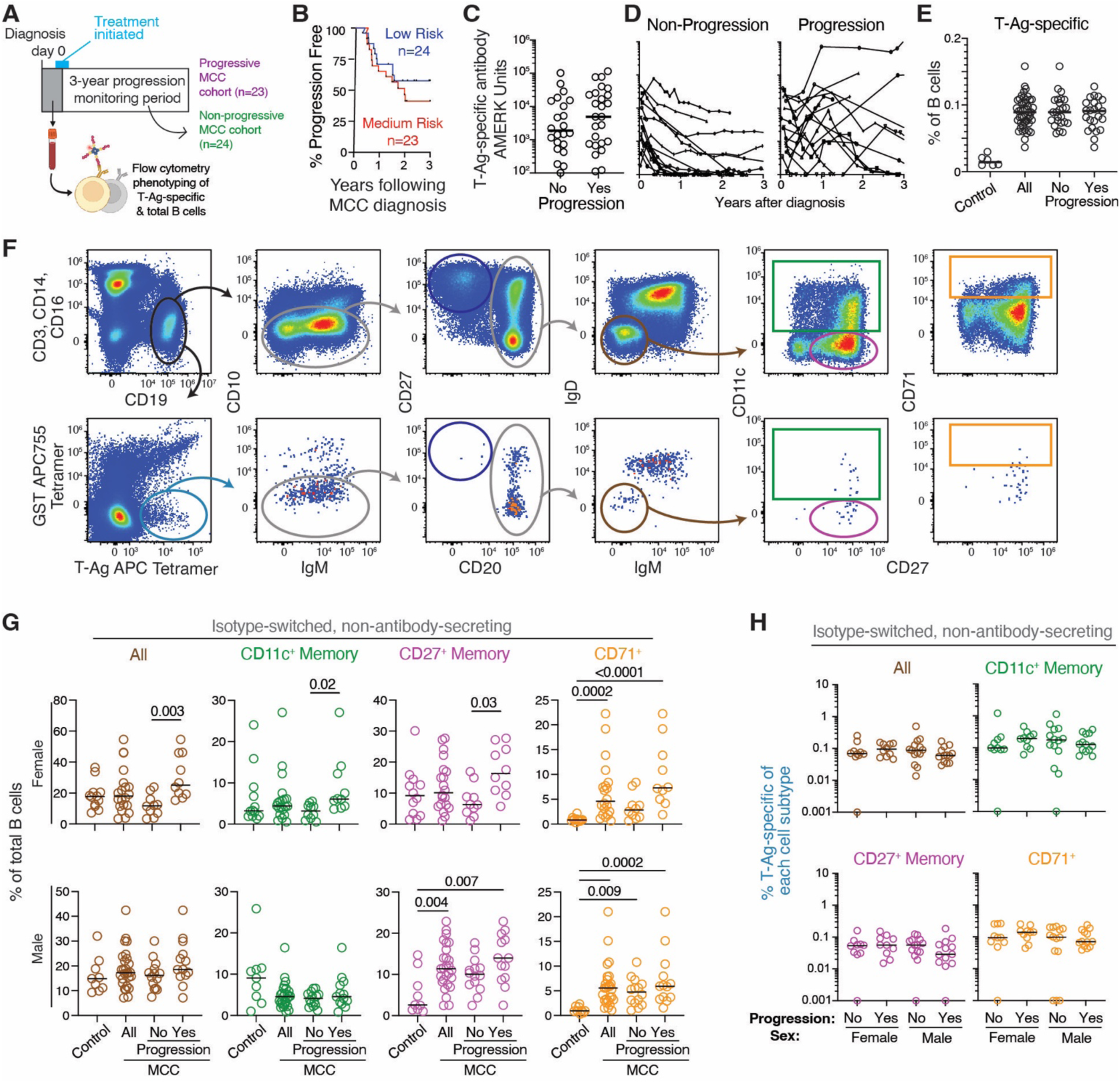
Higher frequency of antigen-experienced B cells in blood of female patients near diagnosis associates with MCC progression. **A**. Schematic representative of prospective cohorts used for experiments analyzing T-Ag-specific antibodies and B cells from MCC patient blood 6-67 days after MCC diagnosis. Patients were monitored for at least 3 years after treatment and stratified into a cohort that experienced progression (n=23) and a cohort that did not experience progression (n=24). **B**. Kaplan-Meier plots displaying percent progression-free survival for patients during the three-year monitoring period following treatment stratified based on clinical risk scoring of data available at the time of MCC diagnosis. **C**. Level of T-Ag-specific antibodies in the blood of MCC patients determined using AMERK near the time of diagnosis for the progression and non-progression cohorts. Bars represent medians. **D**. Longitudinal analysis of T-Ag-specific antibodies in the blood of MCC patients in the progression cohort (n=13) and non-progressive cohort (n=19). Patients for which antibody levels were not available for at least a year after diagnosis were excluded. **E**. Combined data from seven experiments showing the % of B cells binding T-Ag tetramers but not control tetramers in the blood of MCC patient cohorts. Bars represent medians and significant p values determined by non-parametric Dunn’s multiple comparison test are displayed. **F**. Representative flow cytometry of CD27^++^ CD20^−^ CD10^−^ antibody-secreting cells and CD20^+^ IgM^−^ IgD^−^ CD10^−^ isotype-switched B cells within all B cells (top) and T-Ag-specific B cells (bottom) in MCC patient samples. Isotype-switched non-antibody-secreting cells were further subdivided into CD27^+^ memory B cells, CD11c^+^ memory B cells, and CD71^+^ activated B cells. **G**. Pooled data from five experiments showing the % of each cell subtype with the population of total B cells from blood of female (top, n=20) or male (bottom, n=27) patients and age-matched controls with no history of MCC (n=5-6). Data from progressor and non-progressor cohorts is displayed separately and combined. **H**. Pooled data from 4 experiments displaying the % of T-Ag-specific B cells within listed B cell subtype. Bars in **G** and **H** represent medians and p values determined by non-parametric Dunn’s multiple comparison test are displayed when significant.

**Supplemental Figure 3.**
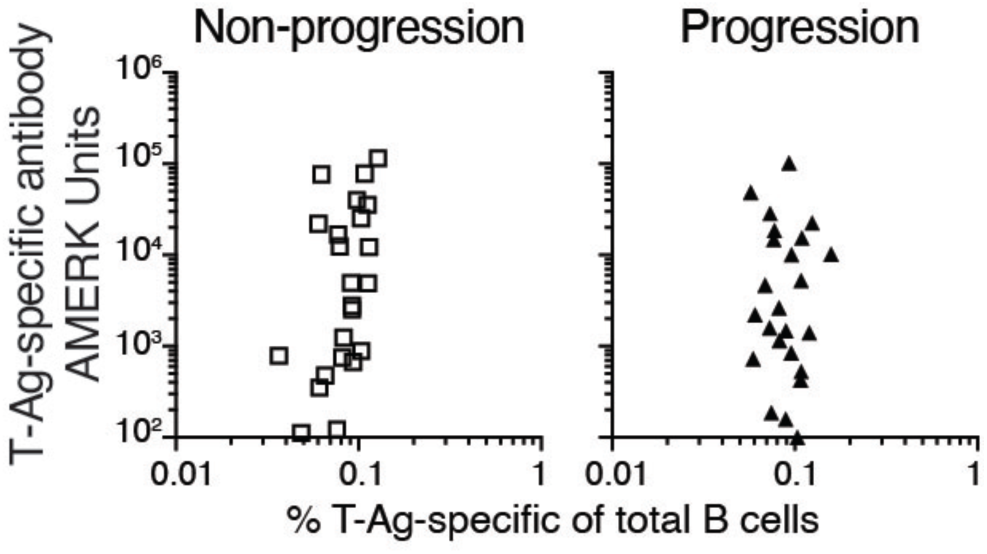
The frequency of T-Ag-specific B cells in the blood does not associate with levels of T-Ag-specific antibodies in the blood or outcome. Analysis of % of B cells specific for T-Ag in the blood versus the level of T-Ag-specific antibodies in the blood prior to treatment for MCC in patients who did not later experience a progression (left) and patients who experienced progression (right).

**Supplemental Figure 4.**
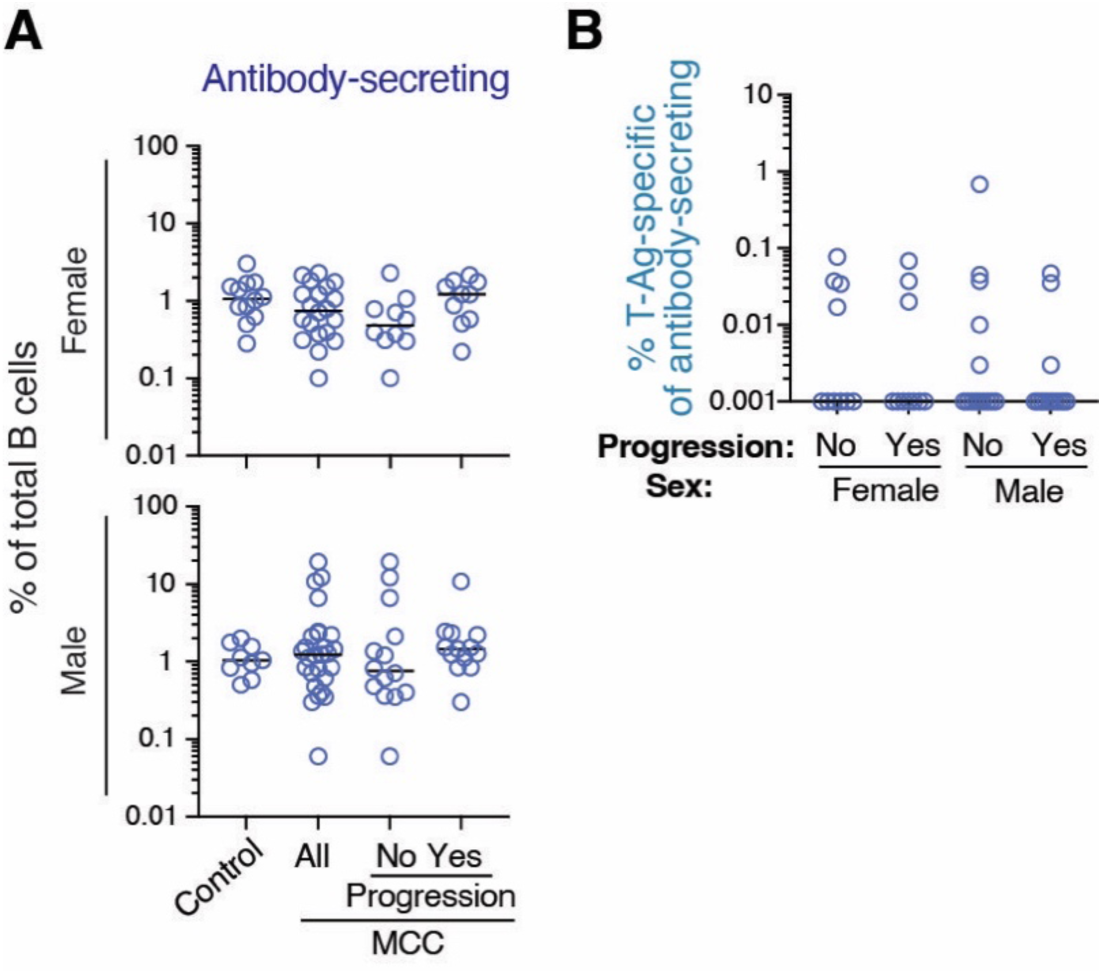
Increases in antibody-secreting cells associates with MCC progression. **A**. Pooled data from four experiments showing the % of CD19^+^ CD27^++^ CD20^−^ CD10^−^ antibody-secreting cells with the population of total B cells (top) or % of T-Ag-specific B cells (bottom) found in PBMC from MCC patients prior to treatment. Data from all patients is displayed combined and stratified based upon whether the patient experienced MCC progression after the analysis time point. **B**. Pooled data from four experiments showing the % of T-Ag-specific cells within the total population of antibody-secreting cells stratified based upon whether the patient experienced MCC progression. Significant p values were generated using a non-parametric Spearman’s correlation test. The bars represent the median and p values calculated using a non-parametric Dunn’s multiple comparison test are displayed when significant.

**Supplemental Figure 5.**
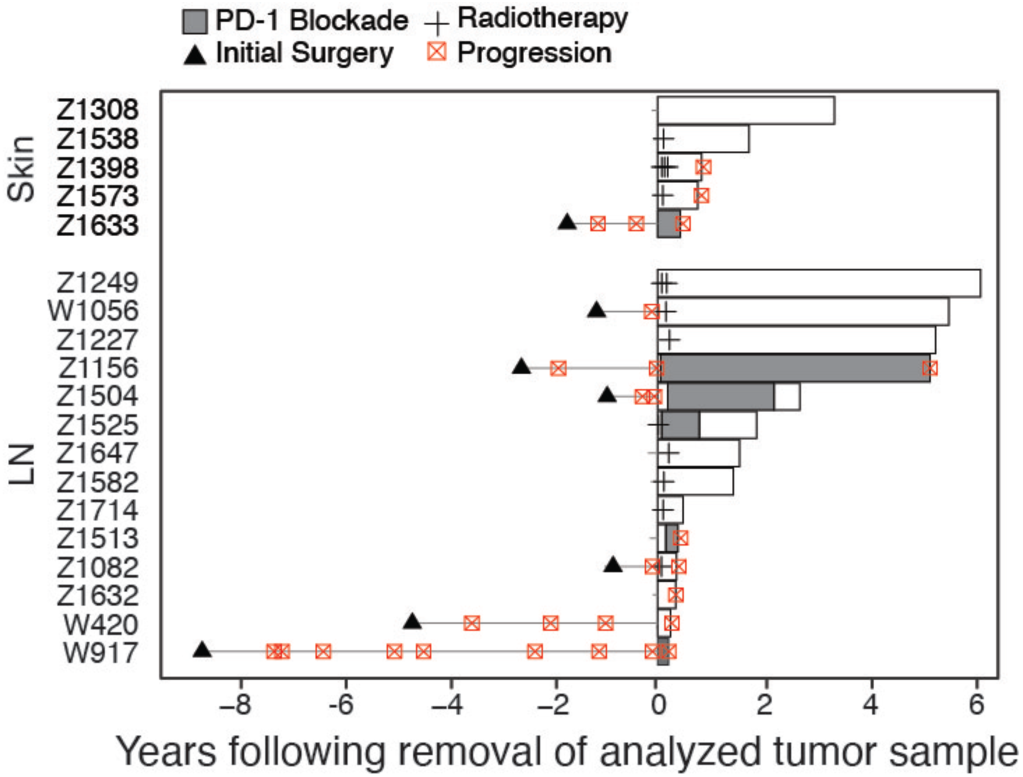
Tumor sample cohort patient history summaries.

**Supplemental Figure 6.**
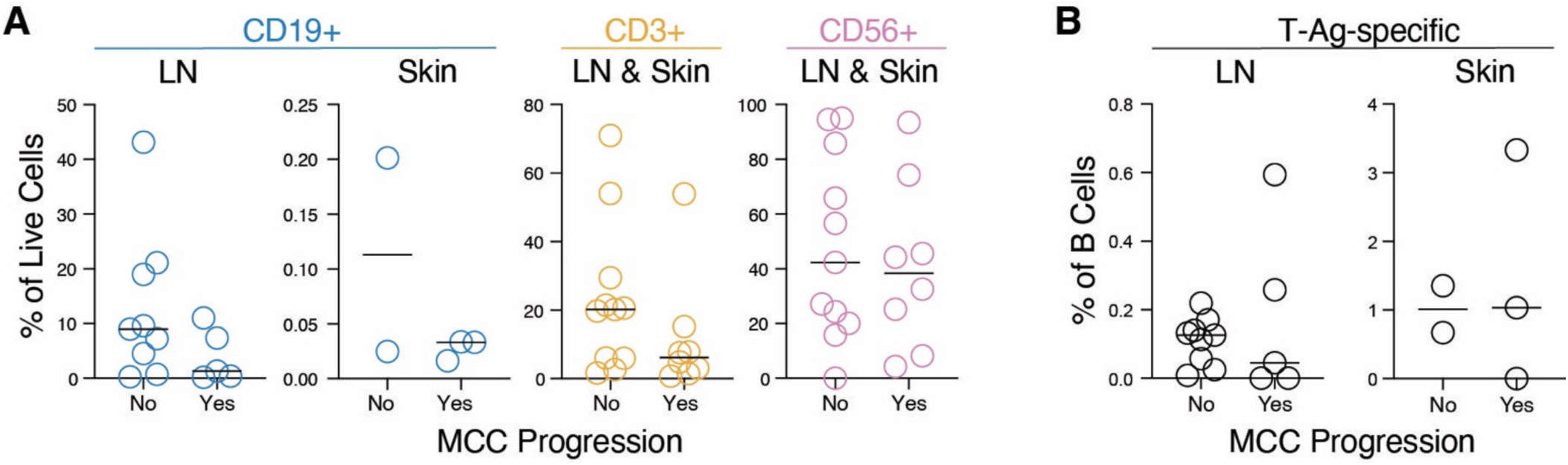
Overall Frequency of T cells, B cells, and MCC cells in tumor samples does not associate with MCC progression after treatment. **A**. Pooled data from five experiments showing displaying the % of live cells that were CD19^+^ B cells, CD3^+^ T cells and CD56^+^ MCC cells found in MCC skin and LN tumor samples using flow cytometry stratified by based on whether the patient experienced MCC progression during the four-year monitoring period. **B**. Pooled data from five experiments showing displaying the % of CD19^+^ B cells that were T-Ag-specific found in MCC skin and LN tumor samples stratified based on whether the patient experienced MCC progression during the four-year monitoring period.

**Supplemental Figure 7.**
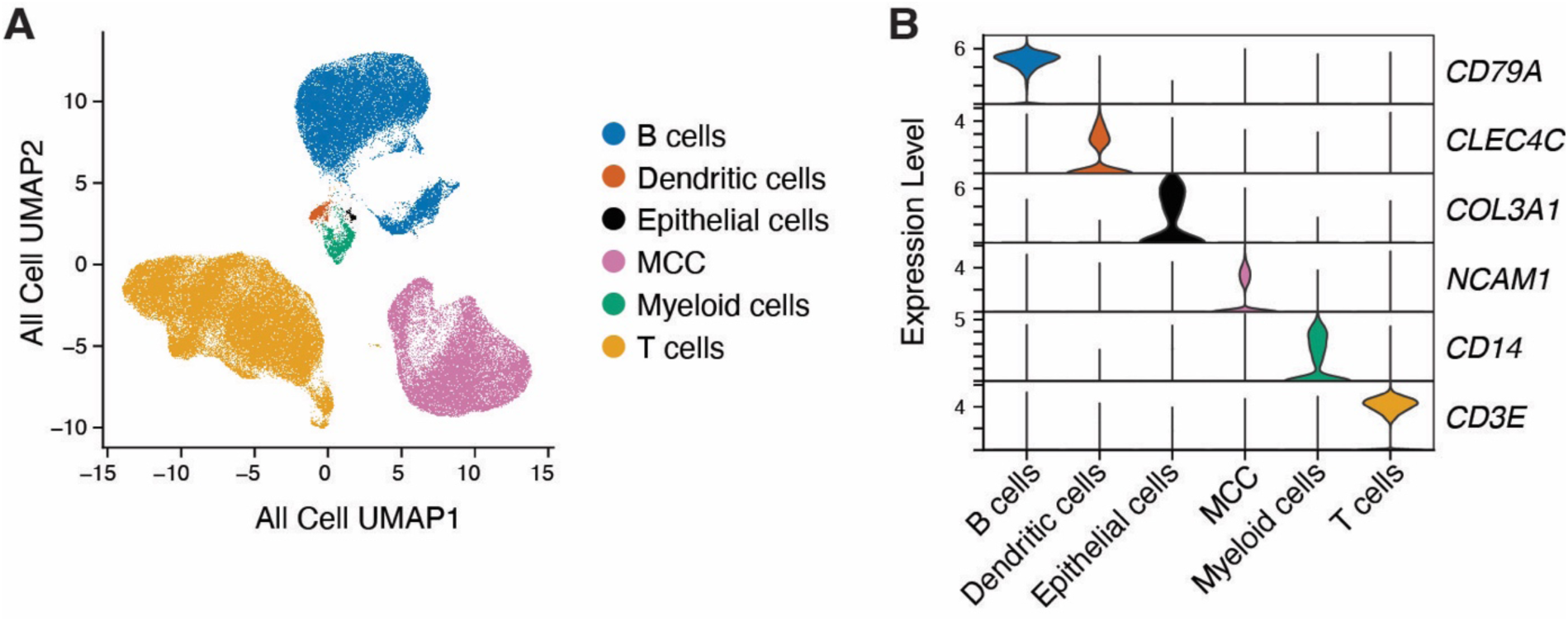
Identification of cell types in UMAP clusters. Expression of key genes used to identify subsets of CD19^+^ B cells, CD3^+^ T cells, CD56^+^ MCC, and a few other contaminating cell types that were not fully excluded during FACS-purification.

**Supplemental Figure 8.**
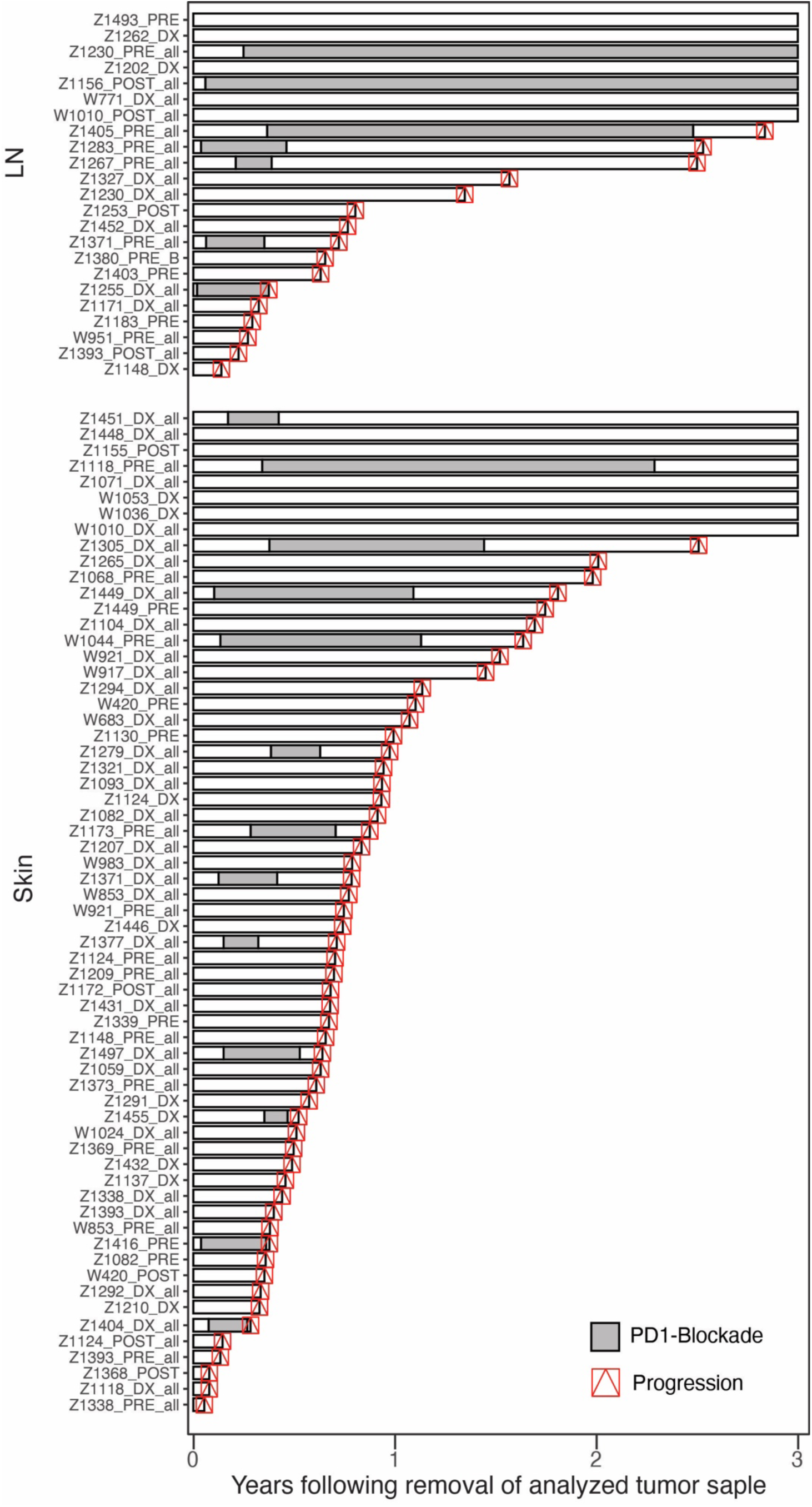
Tumor mIHC cohort patient history summaries.

**Supplemental Figure 9.**
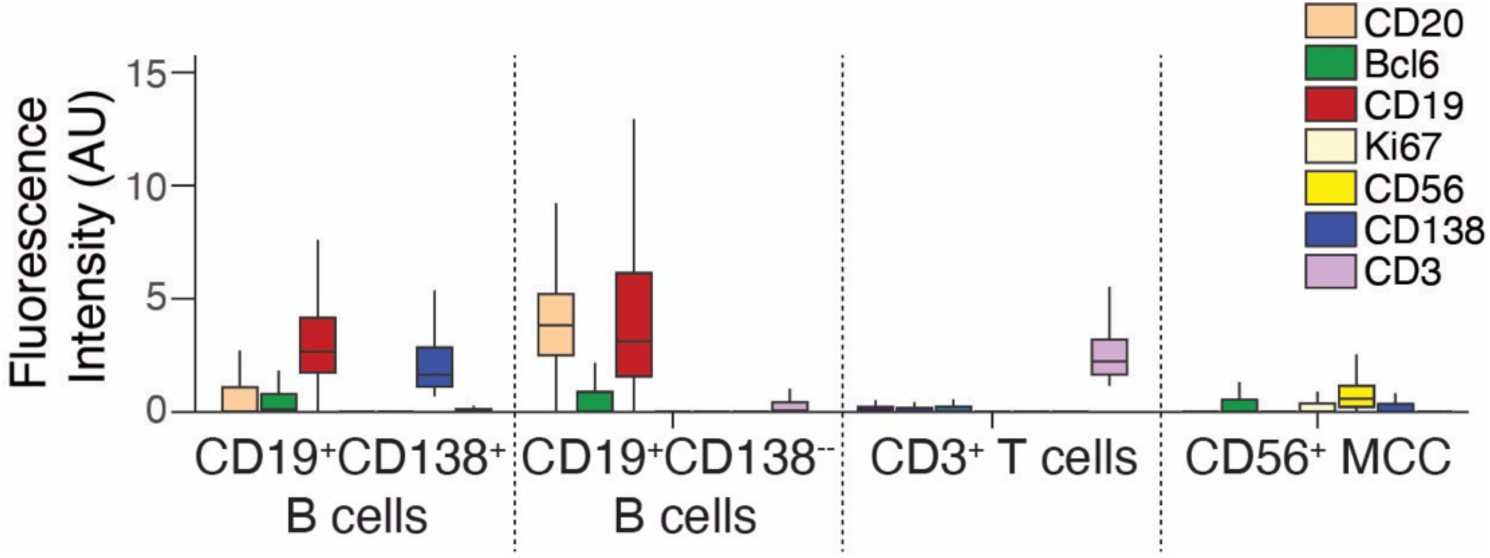
Staining intensity of markers included in the Tumor mIHC panel. Staining intensity of markers used in mIHC experiments are displayed for the identified cell subsets.

**Supplemental Figure 10.**
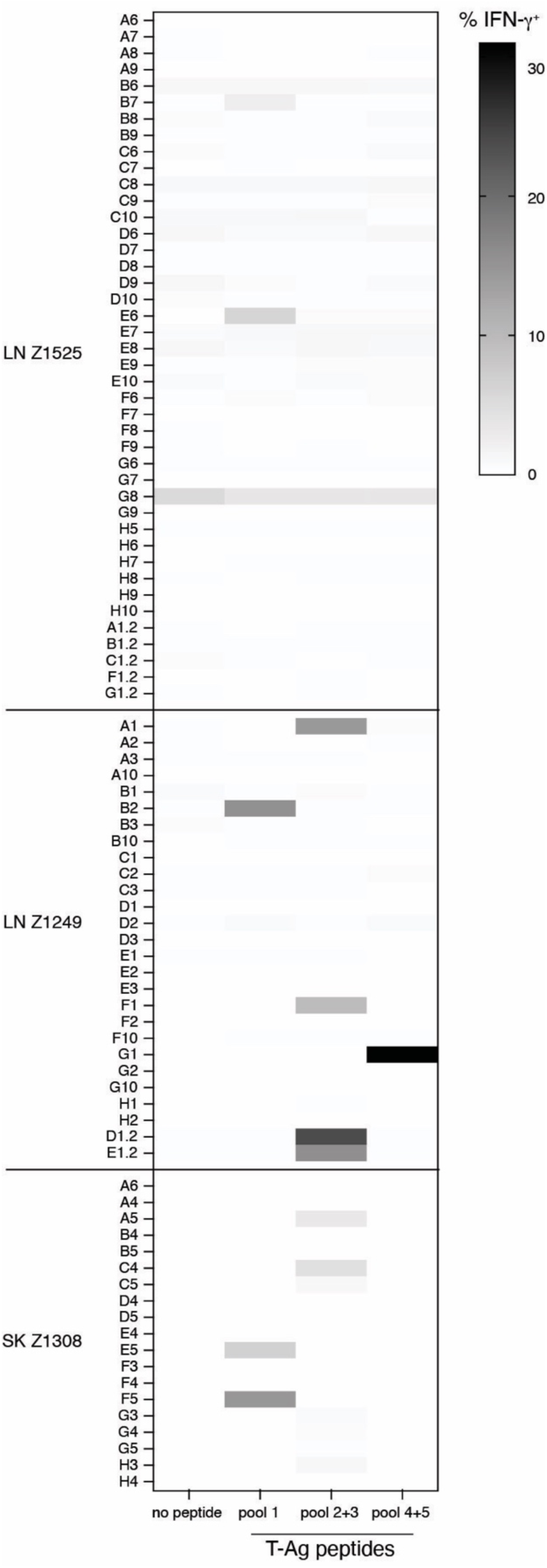
Full TCR Screening. Sixty-nine clonal T_FH_ TCRs from MCC tumor samples were expressed via lentivirus in CD4^+^ cells from blood and incubated with sample-matched B cells in the absence or presence of 0.1 or 5 μg/mL of T-Ag peptide pools for three days prior to assessment of supernatant for secreted IFNγ. Representative of two similar experiments.

## METHODS

### RESOURCE AVAILABILITY

#### Lead contact

Further information and requests for resources and reagents should be directed to and will be fulfilled by the lead contacts, Paul Nghiem and/or Justin Taylor

Materials availability Available upon request.

#### Data and code availability

The expression data obtained in this study has been uploaded to the GEO database with accession no. GSE301498. Any requests for the raw data will be reviewed by the corresponding authors to ensure patient confidentiality is maintained. If possible, the data will be shared under a material transfer agreement. This paper does not report original code, however, code needed to reproduce any of the analyses will be made available upon reasonable request. Any additional information required to reanalyze the data reported in this work is available from the lead contacts upon request.

### EXPERIMENTAL MODEL AND SUBJECT DETAILS

#### Mouse Studies

Animal studies were approved and conducted in accordance with the Fred Hutchinson Cancer Center Institutional Animal Care and Use Committee. Six- to ten-week-old female C57BL/6 mice were obtained from the Jackson Laboratory.

#### Human Studies

Blood samples from MCC patients (**Figure S1, Table S2**) were collected with informed consent for research use and were approved by the Fred Hutchinson Cancer Center (FHCC) institutional review board, in accordance with the Declaration of Helsinki (2013) as part of observational registry studies focusing on Merkel cell carcinoma (IRB protocol 6585). Titers of T-Ag-specific antibodies in blood were determined by the Clinical Immunology Lab at Department of Laboratory Medicine and Pathology at the University of Washington using the AMERK test^13^. Patient samples were selected based on availability of frozen viable PBMCs collected within 67 days of MCC diagnosis. In addition, patient samples were selected to be representative of the sex distribution observed in virally driven MCC. Patient samples were further selected to only include MCPyV-positive cases and those from immune competent patients. Blood samples from age-matched, non-MCC controls were collected with informed consent for research use and were approved by the University of Washington institutional review board, in accordance with the Declaration of Helsinki (2013) (IRB protocol STUDY00001399).

Tumor samples from MCC patients (**Figure S5, S8, Tables S3, S7**) were collected with informed consent for research use and were approved by the Fred Hutchinson Cancer Center (FHCC) institutional review board, in accordance with the Declaration of Helsinki (2013) as part of observational registry studies focusing on Merkel cell carcinoma (IRB protocol 6585). Additionally, tumor samples for single cell experiments (**Figure S5, Table S3**) were selected based on availability of frozen viable tumor digests, number of cells available in sample (>10 million), and included only MCPyV-positive cases. Patient samples were primarily from males due to limited female tumor samples.

Recurrence risk of MCC PBMC patient cohort (**Figure S1, Table S2**) was calculated using an open-source MCC recurrence risk calculator (https://merkelcell.org/prognosis/recur/)^4^. This calculator estimates the probability of recurrence from the time of diagnosis in patients with no prior presentation of MCC. Briefly, percent probability of recurrence for each patient following MCC diagnosis was calculated using known risk factors including age, sex, stage as defined by the American Joint Committee on Cancer, site of primary tumor, and immune status. In contrast, because some MCC patients in single cell RNA-sequencing tumor cohort had prior disease recurrences (**Figure S5, Table S3**), clinical recurrence risk in these patients was independently assessed by 3 blinded clinicians.

Known risk factors used for this assessment included age, sex, MCC stage, presence of metastatic disease, prior recurrences, and immune status. Each patient was then categorized as having “Medium/Low” or “High” probability of MCC recurrence following surgical excision of analyzed tumor.

To assess long-term MCC outcomes in blood studies (**Figure S1, Table S2**), patients were monitored for 3 years starting at the time of biopsy-proven MCC diagnosis. For progression-free survival analyses, an event consisted of a patient having a recurrence or progression of MCC following definitive treatment (surgery, radiation, or anti-PD1 immunotherapy alone or in combination). Patients with no MCC recurrences were censored following 3 years of follow-up.

To assess long-term MCC outcomes in single cell RNA-sequencing tumor studies (**Figure S5, Table S3**) and in tumor imaging studies (**Figure S8, Table S7**), patients were monitored for 4 and 3 years, respectively, starting at the time surgical excision of analyzed tumor. For progression-free survival analyses, an event consisted of a patient having a recurrence or progression of MCC following definitive treatment (surgery, radiation, or anti-PD1 immunotherapy alone or in combination). Patients with no MCC progression were censored following the monitoring period for each study.

### METHOD DETAILS

#### Production of T-Ag, GST, and RSV F proteins

Production of the common T-Ag domain shared by both small and large T-Ag isoforms, was performed as previously described^12^. Briefly, pGEX4T3 plasmids (GE Healthcare) encoding the T-Ag fused to glutathione S-transferase (GST) or GST alone were expressed in Rosetta Escherichia coli and harvested by centrifugation 15 h after induction. Pelleted bacteria were resuspended in 40 mM Tris pH 8.0, 200 mM NaCl, 1 mM EDTA, and 2 mM DTT supplemented with complete protease inhibitor cocktail and lysed by two passes through a Microfluidizer. Lysates were then diluted with an equal volume of glycerol. Pierce Glutathione Spin Columns (ThermoFisher) were used following manufacturer’s instructions to purify GST and T-Ag. After purification, proteins were buffer exchanged by centrifugation using 10kDa Amicon Ultra centrifugal filters into 1xDPBS, aliquoted at 40 μM concentrations, and run on SDS-PAGE to assess protein purification.

Expression plasmid for His-tagged RSV prefusion F antigen was previously described^49^. Briefly, 293F cells were transfected at a density of 10^6^ cells/mL in Freestyle 293 media using 1 mg/mL PEI Max (Polysciences). Transfected cells were cultured for 7 days with gentle shaking at 37 °C. Supernatant was collected by centrifuging cultures at 2500×*g* for 30 minutes followed by filtration through a 0.2 µM filter. The clarified supernatant was incubated with Ni Sepharose beads overnight at 4 °C, followed by washing with wash buffer containing 50 mM Tris, 300 mM NaCl, and 8 mM imidazole. His-tagged protein was eluted with an elution buffer containing 25 mM Tris, 150 mM NaCl, and 500 mM imidazole. The purified protein was run over a 10/300 Superose 6 size exclusion column (GE Life Sciences). Fractions containing the trimeric F protein were pooled and concentrated using 50 kDa Amicon Ultra centrifugal filters (Millipore). The concentrated sample was stored in 50% glycerol at −20 °C.

#### Production of T-Ag and control tetramers

Production of T-Ag, GST, and RSV F tetramers was performed using a previously described protocol^23,24^. Briefly, purified T-Ag, GST, and RSV F proteins were biotinylated using the EZ-Link Sulfo-NHS-LC Biotinylation kit according to manufacturer instructions. Following biotinylation reaction, excess biotin was removed by centrifugation using 10 kDa Amicon Ultra centrifugal filters. Next, the molar ratio of biotin:protein was assessed by Western blot and selected to be ∼1:1 to prevent over- or under-biotinylation of antigen that could reduce the downstream efficacy of protein tetramers. After optimization of biotinylation, T-Ag, GST, and RSV F tetramers were assembled by incubating biotinylated protein with either streptavidin-APC or streptavidin-PE at a 4:1 molar ratio of protein:streptavidin for 30 minutes at room temperature and in the dark. Unconjugated T-Ag and GST were removed by centrifugation using 100 kDa Amicon Ultra centrifugal filters, whereas unconjugated RSV F was removed also by centrifugation but using a 300 K Nanosep centrifugal device (Pall Corporation). Tetramers were buffer exchanged into 1xDPBS by centrifugation using 100 kDa Amicon Ultra centrifugal filter and diluted to 1 μM in 50% glycerol in 1xDPBS for long-term storage at -20C.

To make control tetramers, streptavidin-PE was conjugated to DyLight 594 (PE594) or DyLight 650 (PE650), and streptavidin-APC conjugated to DyLight 755 (APC755) using the DyLight Antibody Labeling kit following manufacturer instructions. After removal of excess dye by centrifugation using 10kDa Amicon Ultra centrifugal filters, biotinylated GST was incubated with streptavidin-PE594 or streptavidin-APC755 at a 4:1 ratio of protein:streptavidin for 30 minutes at room temperature and in the dark. Control tetramers were diluted to 1-3 μM in 50% glycerol diluted with 1xDPBS for long-term storage at -20C.

Oligo-barcoded tetramers were assembled using TotalSeq-C streptavidin-PE. Briefly, biotinylated T-Ag, biotinylated GST-DL594, and biotinylated RSV F were incubated with TotalSeq-C streptavidin-PE containing uniquely identifying oligo tags at a molar 4:1 ratio of protein:streptavidin for 30 minutes at room temperature and in the dark. Unconjugated T-Ag and GST were removed by centrifugation using 100 kDa Amicon Ultra centrifugal filters, whereas unconjugated RSV F was removed also by centrifugation but using a 300 K Nanosep centrifugal device (Pall Corporation).

#### Enrichment and analysis of tetramer-binding mouse B cells

Six- to ten-week-old C57BL/6J female mice were used for experiments. Mice were injected intraperitoneally with 50 μL of CFA emulsion containing 5 μg of T-Ag in 1xDPBS or 1xDPBS alone as control. Seven days post-vaccination, tetramer-binding B cells were enriched using a previously described protocol^23,24^. Briefly, the spleen inguinal, axillary, brachial, cervical, mesenteric, and periaortic lymph nodes for each mouse were pooled and manually dissociated. Following dissociation, 10 mL of DPBS containing 1% heat-inactivated newborn calf serum (sorting buffer) was added, and tissue debris removed, the samples pelleted, and supernatant discarded. For staining of T-Ag-specific B cells, each sample was first incubated with 1 pmol of PE594 control tetramer, 3 pmol of APC755 control tetramer, and 2 μg of anti-Fc receptor antibody 2.4G2 (BioXcell) in 0.2 mL sorting buffer for 10 minutes on ice.

Next, 1 pmol GST APC tetramer, 1 pmol T-Ag PE tetramer, a cell viability dye and fluorescent antibodies against cell-surface lineage markers (CD3, F4/80, Gr-1, B220, CD38, and GL7) were added to samples followed by incubation on ice for 25 minutes. After the incubation, ∼10 mL of sorting buffer was added and the samples centrifuged at 300 × g for five minutes at 4°C and supernatant discarded. Cells were resuspended in 0.15 mL sorting buffer containing 25 μL of anti-APC and 25 μL of anti-PE conjugated microbeads (Miltenyi) and incubated for 30 minutes on ice. Next, 5 mL of sorting buffer was added to each sample and passed over a magnetized LS column (Miltenyi). The tube and column were washed once with 5 mL of sorting buffer and then removed from the magnetic field. Five mL of sorting buffer was pushed through the column with a plunger to elute column-bound cells. Another 5 mL of sorting buffer was plunged through the column to maximize recovery.

For intracellular staining, cells were fixed and permeabilized using the BD Cytofix/Cytoperm buffer according to manufacturer’s instructions. Following permeabilization, an anti-mouse Ig was added and samples incubated for 30 minutes on ice. After incubation, ∼4 mL of sorting buffer was added and the samples centrifuged at 300 × g for five minutes at 4°C and supernatant discarded. Samples were resuspended sorting buffer containing 20,000 AccuCheck Counting Beads (ThermoFisher) analyzed by the BD Fortessa flow cytometer at the Fred Hutch Cancer Center’s Flow Core Facility. UltraComp eBeads Plus (ThermoFisher) or splenocytes were used as single stained controls. Compensation matrix was set up using the BD FACSDiva software. Analysis performed using FlowJo v.10.10.

#### Enrichment and sorting of T-Ag-specific B cells from human blood

Heparinized whole blood from MCC patients was processed at the Specimen Processing Lab (Fred Hutch Cancer Center). PBMC were isolated by routine Ficoll density gradient centrifugation and cryopreserved in freezing medium [50% human serum (Valley Biomedical), 40% RPMI (Corning), and 10% DMSO (Sigma-Aldrich)]. 5-10 × 10^7^ frozen PBMCs from MCC patients or non-MCC controls were thawed into DMEM with 10% fetal calf serum and 100 U/mL penicillin. Cells were centrifuged, resuspended in 100 µL of ice-cold sorting buffer containing 1 pmol of GST PE650 tetramers and 2% rat and mouse serum (ThermoFisher), and incubated at on ice for 10 min. Next, 1 pmol of T-Ag PE tetramers were then added at a final concentration of 5 nM and incubated on ice for 25 min, followed by a 10 mL wash with ice-cold sorting buffer. Next, 25 μL each of anti-APC and anti-PE microbeads (Miltenyi) were added and incubated on ice for 30 minutes, after which 3 mL of sorting buffer was added, and the mixture was passed over a magnetized LS column (Miltenyi). The column was washed once with 5 mL ice-cold sorting buffer and then removed from the magnetic field and 5 mL ice-cold sorting buffer was pushed through the unmagnetized column twice using a plunger to elute the bound cell fraction. Cells were then incubated with 50 μL of sorting buffer containing a cell viability dye and fluorescent antibodies against cell-surface lineage markers (CD3, CD14, CD16, and CD19) for 30 min on ice prior to washing and analysis on a BD FACS Aria sorter at the Fred Hutch Cancer Center’s Flow Core Facility. UltraComp eBeads Plus (ThermoFisher) or PBMCs were used as single stained controls. Compensation matrix and gating of tetramer-binding B cells was set up using the BD FACSDiva software. Tetramer-binding B cells were then individually sorted into empty 96-well PCR plates and immediately frozen.

#### B cell receptor sequencing using single cell RT-PCR

As described previously^50^, individual B cells sorted and frozen into empty 96-well PCR plates, reverse transcription (RT) was directly performed after thawing plates using SuperScript IV (ThermoFisher). Briefly, 3 µL RT reaction mix consisting of 3 µL of 50 µM random hexamers (ThermoFisher), 0.8 µL of 25 mM deoxyribonucleotide triphosphates (dNTPs; ThermoFisher), 1 µL (20 U) SuperScript IV RT, 0.5 µL (20 U) RNaseOUT (ThermoFisher), 0.6 µL of 10% Igepal (Sigma-Aldrich), and 15 µL RNase-free water was added to each well containing a single sorted B cell and incubated at 50°C for 1 h. Following RT, 2 µL of cDNA was added to 19 µL PCR reaction mix so that the final reaction contained 0.2 µL (0.5 U) HotStarTaq Polymerase (Qiagen), 0.075 µL of 50 µM 3′ reverse primers, 0.115 µL of 50 µM 5′ forward primers, 0.24 µL of 25 mM dNTPs, 1.9 µL of 10 × buffer (Qiagen), and 16.5 µL of water. The PCR program was 50 cycles of 94°C for 30 s, 57°C for 30 s, and 72°C for 55 s, followed by 72°C for 10 min for heavy and kappa light chains. The PCR program was 50 cycles of 94°C for 30 s, 60°C for 30 s, and 72°C for 55 s, followed by 72°C for 10 min for lambda light chains.

After the first round of PCR, 2 µL of the PCR product was added to 19 µL of the second-round PCR reaction so that the final reaction contained 0.2 µL (0.5 U) HotStarTaq Polymerase, 0.075 µL of 50 µM 3′ reverse primers, 0.075 µL of 50 µM 5′ forward primers, 0.24 µL of 25 mM dNTPs, 1.9 µL 10 × buffer, and 16.5 µL of water. PCR programs were the same as the first round of PCR. 4 μL of the PCR product was run on an agarose gel to confirm the presence of a ∼500-bp heavy chain band or ∼450-bp light chain band. 5 μL from the PCR reactions showing the presence of heavy or light chain amplicons was mixed with 2 µL of ExoSAP-IT (ThermoFisher) and incubated at 37°C for 15 min followed by 80 °C for 15 min to hydrolyze excess primers and nucleotides. Hydrolyzed second-round PCR products were sequenced by Genewiz with the respective reverse primer used in the second-round PCR, and sequences were analyzed using IMGT/V-Quest to identify V, D, and J gene segments. Paired heavy chain *VDJ* and light chain *VJ* sequences were cloned into pTT3-derived expression vectors containing the human IgG1, IgK, or IgL constant regions using In-Fusion cloning (Clontech).

#### Monoclonal antibody production

Secreted T-Ag and control IgG1 antibodies were produced by Genscript, WuXi, or in house as described previously^50^. For the latter, 10^6^ cells/mL of 293F cells were co-transfected with heavy and light chain expression plasmids at a ratio of 1:1 in Freestyle 293 media (ThermoFisher) using 1 mg/mL PEI Max. Transfected cells were cultured for 7 days with gentle shaking at 37 °C. Supernatant was collected by centrifuging cultures at 2500×*g* for 15 minutes followed by filtration through a 0.2 µM filter. Clarified supernatants were then incubated with Protein A agarose (ThermoFisher) followed by washing with IgG-binding buffer (ThermoFisher). Antibodies were eluted with IgG Elution Buffer (ThermoFisher) into a neutralization buffer containing 1 M Tris-base pH 9.0. Purified antibody was concentrated and buffer exchanged into 1xDPBS using 50 kDa Amicon Ultra centrifugal filters.

#### Bio-Layer Interferometry

Bio-layer interferometry (BLI) assays were performed on the Octet.Red instrument (ForteBio) at room temperature with shaking at 500 rpm. To assess binding of monoclonal antibodies to T-Ag, streptavidin capture sensors (ForteBio) were loaded in kinetics buffer (PBS with 0.01% bovine serum albumin, 0.02% Tween 20, and 0.005% NaN_3_, pH 7.4) containing 1 µM biotinylated GST as control or biotinylated T-Ag for 2.5 minutes. After loading, the baseline signal was recorded for 1 minute in a kinetics buffer. The sensors were then immersed in kinetics buffer containing 40 µg/mL of purified monoclonal antibody for a five-minute association step followed by immersion in kinetics buffer for a five-minute dissociation phase. The binding curves were generated after subtracting the background signal from each analyte-containing well using a negative control mAb at each time point. Curve fitting was performed using a 1:1 binding model and ForteBio Octet data analysis software release 9.0.

#### Spectral flow cytometry of B cells in MCC patient blood

PBMC from MCC patients (**Figure S2, Table S2)** were analyzed by spectral flow cytometry. Briefly, frozen PBMC were thawed into DMEM with 10% fetal calf serum and 100 U/mL penicillin. Cells were centrifuged and resuspended in 50 µL of ice-cold sorting buffer. GST APC755 was added at a final concentration of 1.25 pmol in the presence of 2% rat and mouse serum (ThermoFisher) and incubated at room temperature for 10 min. Cells were then incubated with 50 µL of sorting buffer containing a cell viability dye, 0.5 pmol of T-Ag APC tetramer, and fluorescent antibodies against cell-surface lineage markers (CD3, CD14, CD16, CD19, CD20, CD10, CD27, CD11c, CD71, IgM, IgD, and IgG) for 30 min on ice prior to washing and analysis on a Cytek Aurora spectral analyzer at the University of Washington, Department of Immunology’s Cell Analysis Facility. Antibody capture beads (ThermoFisher) or cells were used to compensate each fluorophore in the experiment. Spectral unmixing was performed using SpectroFlo software. Analysis performed using FlowJo v.10.10.

#### Tumor single cell RNAseq sample preparation

Fresh MCC tumor specimens from needle cores, punch biopsies, or surgical excisions were processed into single-cell digests by mincing them into small pieces with sterile forceps and scissors, followed by incubation in 20 mL of digestion medium composed of RPMI plus 0.002 g DNase (Worthington Biochemical), 0.008 g collagenase (Worthington Biochemical), and 0.002 g hyaluronidase (Worthington Biochemical) in a 10-cm dish at 37°C with frequent, gentle swirling. After 3 hours of digestion, cells were strained through a 70 μm filter, centrifuged, resuspended in freezing medium and stored in liquid nitrogen.

Frozen cells were thawed at 37°C, followed by dropwise addition of 1 mL complete media (RPMI, 10% Fetal bovine serum, 100 U/mL penicillin, 100 μg/mL streptomycin). Four Equivolume additions of complete media was added dropwise with gentle mixing in between additions for a total volume of 32 mL. Thawed cells were washed twice with 4°C 1xDPBS and transferred to 5 mL FACS tubes (ThermoFisher). Live/dead stain was added, followed by a blocking buffer to bring samples to 0.5% BSA, 5% TruStain FcX buffer (BioLegend), 100 nM dasatinib, and control tetramers (oligo-PE-GSTDL594 and APCDL755-GST). Samples were then incubated on ice for 10 min followed by the following reagents in order: tetramers (T-Ag PE-oligo tetramer, T-Ag APC tetramer, RSV F PE-oligo tetramer, and RSV F APC tetramer), oligo-hashtag antibodies to identify sample origin in subsequent pooling steps, fluorochrome-labeled antibodies, and oligo-labeled antibodies. Cells were then incubated on ice for 30 min and washed three times with sorting buffer. Cells were then sorted on an Aria II Cell sorter (BD Biosciences). After exclusion of cell debris, dead cells, PE/APC tetramer-binding CD19^+^ CD3^−^ CD56^−^ PE594^−^ APC755^−^ B cells, tetramer-negative B cells CD19^+^ CD3^−^ CD56^−^ PE594^−^ APC755^−^ B cells, CD3^+^ CD19^−^ CD56^−^ T cells, and CD56^+^ CD3^−^ CD19^−^ cells were sorted into cold complete media, pooled and immediately prepared for sequencing. For most experiments, tetramer^+^ CD19^+^ B cells were sorted into a tube with CD3^+^ T cells and kept separate from Tetramer^−^ CD19^+^ B cells that were sorted into a tube with CD56^+^ MCC cells.

#### Single cell RNA-sequencing

Single cell suspensions sorted from tumors and brought to a concentration of 700-1,200 cells/mL, loaded into microfluidic chip G (10X Genomics), and run through a Chromium controller to obtain Gel Beads-in-Emulsion (10X Genomics). Resulting cell suspensions then went through a library preparation process for single-cell RNA-sequencing along with paired V(D)J-seq for BCR and TCR receptor clonotypes using the 5’ transcriptome kit with feature barcoding (V2; 10X Genomics) per manufacturer guidelines. The cDNA library was enriched for 1056 human immunology panel genes (10X Genomics) and sequenced using a NovaSeq instrument (Illumina) with 2 x 92 base pair paired-end reads aiming for an average of 20,000 reads/cell.

#### Single-cell RNA-sequencing data analysis

Raw sequencing reads were aligned to the hg38 genome using Cell Ranger v.8.0.1. Filtered counts matrices of transcripts and feature barcoding counts were loaded into a SingleCellExperiment object for further analyses in R (v.4.1.2). Sample hash deconvolution was performed using DropletUtils (v.1.14.2). Doublets were detected using scds (v.1.10.0) and hash deconvolution and subsequently removed.

Cells from different runs were then integrated using the mutual nearest neighbor method though the batchelor package (v1.10.0). UMAP dimensionality reduction was performed using the integrated values. Clustering was performed using the integrated transcript values reads through the walktrap algorithm on a nearest neighbor graph (scran v.1.22.1). Numbers of clusters was varied by scaling the number of nearest neighbors (k) during graph construction followed by analysis via clustree (v.0.5.0). Clusters were then labeled as major cell lineages of T cells, B cells, myeloid cells, epithelial cells, dendritic cells, and tumor cells through expression of key genes including *CD79A, CLEC4C, COL3A1, NCAM1, CD14,* and *CD3E*. Cluster labels were then validated by investigating the portion of cluster with productive BCR or TCR rearrangements. Cell lineages were then isolated *in silico* and dimensionality reduction and B cells re-clustered as above. Clusters were labeled based on expression of lineage markers (**Figure 3b**).

To identify T-Ag-specific B cells in tumors, we analyzed T-Ag oligo reads and control antigen oligo reads in B cells. T-Ag-specific CD19^+^ B cells were defined as cells with T-Ag oligo reads at least one log higher than background and fewer than 20 reads from GST or RSV F control antigens. Additionally, eight B cell families in which fewer than 50% of clones met this threshold were classified as non-specific. Sequences from tested antibodies were obtained from BCR sequencing analyses and are reported in **Table S10**.

#### BCR sequencing analysis

Raw output files were demultiplexed and processed using CellRanger v.8.0.1 software (10X Genomics). For each donor, publicly available reference sequences from the IMGT/V-QUEST reference directory at https://www.imgt.org/ were used to identify V, D, and J gene segments. Next, data were processed and analyzed using the Immcantation Framework (http://immcantation.org) with Change-O v.1.0.2. For each 10X dataset, the filtered_contig.fasta file was annotated using IgBlast v.1.16 with the related donor-specific VJ genes database. To generate adaptive immune receptor repertoire (AIRR) rearrangement data, the filtered_contig_annotations.csv file was used, and only productive sequences were kept. The heavy and light chain sequences were separated in two files. The threshold for trimming the hierarchical clustering of B cell clones was determined by the SHazaM module for determining distance to nearest neighbor. With the Change-O DefineClones function, clones were assigned based on IGHV genes, IGHJ gene and junction distance calculated by SHazaM (distance 0.14). The generated clone-pass file was verified and corrected using the Change-O light_cluster function, based on the analysis of the light chain partners associated with the heavy chain clone. Independent clone-pass files were generated for each 10X run. For downstream analysis, all clone-pass files from were combined and re-clustered all together. Germlines were reconstructed using the Change-O CreateGermlines function. To obtain the final AIRR format file containing paired information on the same row, we used a Java script to process and filter the sequences. Only the heavy chains paired with one κ and/or one λ were filtered in for downstream analysis. Number and frequency of IgH mutations for each b cell were obtained using the SHazaM observedMutations function. Clonal families were visualized using the R package ggalluvial.

#### Multiplex immunohistochemistry of MCC tumors

Formalin-fixed paraffin-embedded tissues were stained on a Leica BOND Rx autostainer using the Akoya Opal Multiplex IHC assay (Akoya Biosciences) with the following changes: Additional high stringency washes were performed after the secondary antibody and Opal fluor applications using high-salt TBST (0.05M Tris, 0.3M NaCl, and 0.1% Tween-20, pH 7.2-7.6). TCT was used as the blocking buffer (0.05M Tris, 0.15M NaCl, 0.25% Casein, 0.1% Tween 20, pH 7.6 +/- 0.1). Primary antibodies were incubated for 1 hour at room temperature. Tissues were counterstained with DAPI to identify the nuclei.

Slides were mounted with ProLong Gold and cured for 24 hours at room temperature in the dark before image acquisition at 20x magnification on the Akoya PhenoImager HT Automated Imaging System. Images were spectrally unmixed using Akoya inForm software.

#### Multiplex immunohistochemistry image analysis

HALO software (Indica Labs) High-Plex FL module was used to analyze each TMA slide image. Annotations of tumor and stroma sections were performed by a licensed pathologist. Cells were then identified based on the DAPI nuclear stain, and mean pixel fluorescence intensity was measured in applicable compartments for each cell. To account for variability across the TMA, cell detection algorithms were optimized individually for each TMA core to ensure accuracy. While many cores ultimately shared the same algorithm parameters, each core was initially assessed independently to determine the most appropriate settings using real-time tuning. The final algorithm parameters for each core were then applied accordingly. Two layers of quality control were performed: 1) comparing positive cell signals to positive object data results on at least six different cores, and 2) comparing cell counts to slide images reviewed by a pathologist. Summary data were exported to CSV files, and cell counts were merged with corresponding patient metadata using STATA 16. GraphPad PRISM 9 and R version 4.2.1 (R Foundation for Statistical Computing) were used to generate plots and compare patient treatment groups.

#### TCR analysis and lentiviral expression in primary CD4^+^ T cells

TCR analysis was performed using some functions from Scirpy to assess clonal expansion and calculate clonotype size as previously described^37^. For TCR construction, the PRRL vector was modified with six start codon a promoter point mutations and cysteines were introduced to mediate TCR pairing as previously described^37^. TCR constructs were synthesized and sequence verified (Twist) containing codon-optimized DNA fragments of *TRBV*-*CDR3*-TRBJ-*TRBC* followed by a P2A skip sequence, TRAV-CDR3-TRAJ-TRAC cloned into PRRL-SIN. Lentivirus was produced freshly for each experiment. For this, Lenti-X cells (Clontech) were transiently transfected with each TCR vector and psPAX2 (Addgene plasmid no. 12260) and pCMV-VSV-G (Addgene plasmid no. 8454) packaging plasmids. Two days later, lentiviral supernatant was harvested from Lenti-X cultures, filtered using 0.45-mm polyethersulfone (PES) syringe filters (Millipore), and added to stimulated CD4^+^ T cells in a 48-well tissue culture plate.

For T cell transduction, CD4^+^ T cells from control PBMC were thawed and stimulated with anti-CD3/anti-CD28 microbeads at a 3:1 bead:cell ratio (Dynabeads, Invitrogen) in T cell media containing L-glutamine and HEPES (Gibco) supplemented with 10% human serum, 50 mM beta-mercaptoethanol, penicillin, streptomycin, 4 mM L-glutamine, 50 U/mL IL-2, and 5 ng/mL IL-7 in RPMI for 2 days. On day 2, magnetic beads were removed, and cells were nucleofected using a Lonza 4D nucleofector in 100 mL of buffer P3 using program EH-115. TCR was knocked out using CRISPR targeting the first exon of the *TRBC* and *TRAC* alone as previously described^37^. For this, 80 mmol/L *TRAC* RNA (IDT), 80 mmol/L *TRBC* RNA (IDT), and 80 mmol/L of the gRNA AGAGTCTCTCAGCTGGTACA in duplex buffer (IDT), were mixed well and incubating in a heating block at 95°C for 5 minutes followed by slowly cooling to room temperature. The resulting 40 mmol/L duplexed RNA was mixed with an equal volume of 24 mmol/L Cas9 (IDT) and 1/20th volume of 400 mmol/L Cas9 electroporation enhancer (IDT) and incubated at room temperature for 15 minutes prior to nucleofection. Cells were allowed to rest for 4 hours in media prior to lentiviral transduction. Polybrene (Millipore) was added to a final concentration of 4.4 mg/mL, and cells were centrifuged at 800 x *g* and 32°C for 90 minutes. Sixteen hours later, viral supernatant was replaced with fresh T cell media. Half of the media was removed and replaced with fresh T cell media every 48 to 72 hours. Transduction efficiency into TCR knockout T cells was determined one week after nucleofection by measuring CD3 expression in each transduced sample compared to TCR knockout control T cells.

#### CRISPR engineering of B cells

B cells were engineered to express T-Ag-specific antibodies as described previously^38^. Briefly, human B cell medium was IMDM supplemented with 10% FBS (Gemini Biosciences or Peak serum), 100 U/mL penicillin and 100 μg/mL streptomycin (Gibco), except in antibiotic free steps as noted. Blood was obtained from adult volunteers by venipuncture and was approved by the Fred Hutch Institutional Review Board. Informed consent was obtained before enrollment. PBMCs were isolated from whole blood using Accuspin System Histopaque-1077 (Sigma-Aldrich) resuspended in 10% dimethylsulfoxide in heat-inactivated fetal bovine serum and cryopreserved in liquid nitrogen before use. PBMCs were thawed and B cells isolated using negative selection using the Human B Cell Isolation Kit II (Miltenyi) according to the manufacturer’s recommendations. Isolated B cells were resuspended at 0.5-1.0×10^6^ cells/mL in stimulation media, which consisted of human B cell medium supplemented with 100 ng/mL MEGACD40L (Enzo Life Sciences), 50 ng/mL recombinant IL-2 (BioLegend), 50 ng/mL IL-10 (Shenandoah Biotech), 10 ng/mL IL-15 (Shenandoah Biotech), 1 μg/mL CpG ODN 2006 (IDT) and incubated . After 48 hours, cells were electroporated using the Neon Transfection System. Cas9 (Invitrogen) and gRNA (Synthego) were precomplexed at a 1 to 2 molar ratio in Neon Buffer T for 20 minutes at room temperature. Cells were washed with 1xDPBS and resuspended to 2.5×10^7^ cells/mL in Neon Buffer T containing 12 μg of precomplexed gRNA/Cas9 per 10^6^ cells. The cell/huIgH_296_ gRNA/Cas9 mixture was electroporated with one 20 millisecond pulse at 1750V and immediately plated into stimulation media as described above, without antibiotics. After 30 minutes, AAV was added to a final concentration of up to 20% culture volume amounting to a MOI of 10^5^-10^6^ genome copies per cell and incubated for 2 - 4 hours. AAVs were produced by VectorBuilder containing an equimolar concentration of antibody constructs corresponding to 1G04, 2H04, and 1B09 from containing the full antibody light chains physically linked to the heavy chain VDJs with a linker^39^ containing Strep-tagII^40–42^. Cells were next transferred to a larger culture dish to allow for further expansion for two days. For secondary expansion, every four days B cells were passaged onto irradiated (80 gy) NIH 3T3-CD40L feeder cells in human B cell medium containing 5 μg/mL human recombinant insulin (Sigma), 50 μg/mL transferrin (Sigma), 50 ng/mL human IL-2 (BioLegend), 20 ng/mL human IL-21 (BioLegend), and 10 ng/mL human IL-15 (Shenandoah Biotech).

#### Stimulation of T-Ag-specific CD4^+^ T cells by B cells

Engineered or control B cells from the same donor as T cells were isolated from thawed PBMC by positive selection using human CD19 microbeads (Miltenyi) according to the manufacturer’s instructions. Isolated B cells were cultured with irradiated (5000Gy) NIH 3T3 cells expressing human CD40L in B cell medium supplemented with 200 U/ml human IL-4 (Peprotech). B cells were harvested and restimulated with 3T3 CD40L every 7 days and supplemented with fresh medium and cytokines every 3 days. For tumor lysate preparation, WaGa tumor cells were counted and resuspended at 5×10^6^/mL in serum free RPMI media. Cell suspension was frozen/thawed 5 times alternating between three minutes submerged in liquid nitrogen and three minutes in a 37 °C water bath. Solution was then centrifuged for 5 minutes at 1000 x G to remove debris and remaining cells. Supernatant was then transferred to a fresh tube to be used as tumor lysate. 25 µL lysate was added to co-cultures for a total volume of 200µL per well. 10^5^ B cells were mixed with 50,000 transduced T cells in the presence or absence of 0.1 or 5 mg/mL of T-Ag peptide pools, 0.1 - 10 mg/mL of purified T-Ag, or crude lysate derived from WaGa MCC cells for 24 hours prior to detection of IFNγ in culture supernatant using the using an ELISA kit (ThermoFisher). Alternatively, the expression of OX40 and CD40L was measured using flow cytometry.

#### Quantification and statistical analysis

The statistical tests applied were two-sided unless specified otherwise. T tests were used to compare differences between two groups unless otherwise noted. When comparing more than two groups, the nonparametric Kruskal–Wallis test was used. Fisher’s exact test was used to evaluate differences between two categorical variables. The significance levels for Kaplan–Meier analyses were calculated using a two-sided log rank test. GraphPad Prism 9 and R version 4.2+ (R Foundation for Statistical Computing) were used to generate plots and compare patient treatment groups.

## Notes

**Conflicts of Interest:** J.J.T, P.N., D.M.K., D.A.G. and H.J.R. are co-inventors on institutionally owned patent application related to this work. J.J.T. is an advisor for Bespoke Biotherapeutics. P.N.’s institution has received grant support from EMD Serono and Bristol Myers Squibb as well as honoraria from Merck and EMD-Serono unrelated to this work. P.N. is a co-inventor on institutionally owned patents concerning MCC but unrelated to this work. J.J.T. is co-inventor on institutionally owned patents unrelated to this work. J.J.T. has received research funding from Vir Biotechnology, Merck & Co, and IGM Biosciences and honoraria from AstraZeneca and Genentech that is unrelated to this work.

### Competing Interest Statement

J.J.T, P.N., D.M.K., D.A.G. and H.J.R. are co-inventors on institutionally owned patent application related to this work. J.J.T. is an advisor for Bespoke Biotherapeutics. P.N.s institution has received grant support from EMD Serono and Bristol Myers Squibb as well as honoraria from Merck and EMD-Serono unrelated to this work. P.N. is a co-inventor on institutionally owned patents concerning MCC but unrelated to this work. J.J.T. is co-inventor on institutionally owned patents unrelated to this work. J.J.T. has received research funding from Vir Biotechnology, Merck and Co, and IGM Biosciences and honoraria from AstraZeneca and Genentech that is unrelated to this work.

### Summary of Updates

Title revised and conflict of interest added to the first page

### Funding

- National Cancer Institute: P01 CA225517

